# A mechanistic mathematical model of initiation and malignant transformation in sporadic vestibular schwannoma

**DOI:** 10.1101/2021.10.03.457528

**Authors:** Chay Paterson, Ivana Bozic, Miriam J. Smith, Xanthe Hoad, D. Gareth R. Evans

## Abstract

A vestibular schwannoma (VS) is a relatively rare, benign tumour of the eighth cranial nerve, often involving alterations to the gene *NF2*. Previous mathematical models of schwannoma incidence have not attempted to account for alterations in specific genes, and could not distinguish between point mutations and loss of heterozygosity (LOH). Here, we present a mechanistic approach to modelling initiation and malignant transformation in Schwannoma. Each parameter is associated with a specific gene or mechanism operative in Schwann cells, and can be determined by combining incidence data with empirical frequencies of pathogenic variants and LOH. This results in new estimates for the base pair mutation rate *u* = 4.48 × 10^−10^ and the rate of LOH = 2.03 × 10^−6^/yr in Schwann cells. In addition to new parameter estimates, we extend the approach to estimate the risk of both spontaneous and radiation-induced malignant transformation. We conclude that radiotherapy is likely to have a negligible excess risk of malignancy for sporadic VS, with a possible exception of rapidly growing tumours.

## 1 Introduction

Vestibular schwannoma is a benign tumour of Schwann cells on the eighth cranial nerve. It is a relatively rare disease, with a lifetime risk of around 1 in 1000 [1, 2]. The majority of vestibular schwannoma cases carry mutations on the gene *NF2* and at least one other hit [3]. Age-related risk is described relatively well by a multi-stage model involving three alterations [4]. This is consistent with a picture in which most cases involve two hits to *NF2*, and one additional insult [3, 5].

Vestibular schwannomas are usually benign, very rarely undergoing malignant transformation: only around 0.2% of cases develop into malignancy [2]. A recent study on familial neurofibromatosis type-II patients (in which *NF2* loss is inherited) suggested an association between radiotherapy and malignant transformation [6]. From such a small sample size, it was impossible to give a quantitative estimate for the excess risk associated with irradiation [7].

Due to the rarity of malignant schwannoma, its pathogenesis is unclear, and the relationships between genomic features, clinical course, and epidemiology are poorly understood [5]. The benign disease is better understood: researchers have been able to use sequencing studies to investigate what genetic alterations have occurred in individual cases [3, 8, 9]. A mathematical model that could connect genomic data to clinical observables would therefore be very useful. Proposed models should pay close attention to specific, plausible mechanisms; as well as summarising the experimental literature in a precise, quantitative form.

One recent example of a mechanistic model connects every parameter to either a population of cells, or the rate of a specific biological process: for example, the mutation rate of individual genes, or the rate of cell division in precursor cells [10]. Several key parameters can be determined directly from the sequences of implicated genes, with no statistical fitting. The main result is an incidence curve for a specific molecular subtype involving alterations to three genes, with the additional advantage that every parameter has a clear interpretation in terms of a specific mechanism [4, 10, 11].

Here, we extend this approach to meet three main goals: first, to describe the incidence of sporadic vestibular schwannoma with a mechanistic model. In this process, we find a new use for existing experimental data, using it in section 2.1 to derive new estimates for biological parameters of interest. Secondly, to use these parameter estimates in a new model for the lifetime risk of malignant transformation. Our work in section 3 may help to constrain the genes responsible. Thirdly, to model the excess risk of malignancy following a dose of ionizing radiation. At each stage, by relating model parameters to potential observables like variant allele frequencies, we are able draw new inferences from existing data, and suggest informative future experiments.

## 2 Modelling incidence of sporadic vestibular schwannoma

The incidence of sporadic vestibular schwannoma is relatively well-described by a three hit model [4]. Our model is also a three hit process, but is distinguished from previous efforts by a focus on specific genes; by allowing for hits to occur in any order; and by tying all parameters to underlying mechanisms. For simplicity, we will not study familial or germline *NF2* mutations, and focus only on sporadic VS involving somatic *NF2* mutations.

*NF2* displays somatic mutations in 85% of all sporadic vestibular schwannomas according to current estimates, a majority [3]. To reduce the number of unknowns, we consider *only* cases of VS associated with somatic *NF2* loss. In addition to point mutations on *NF2*, loss of heterozygosity (LOH) is also commonly found on chromosome 22 [3, 8]. *NF2* loss can only account for two hits: at least one other gene must be involved. A natural hypothesis is that the third hit is an oncogene (or one of several possible proto-oncogenes). However, the third hit may also be a tumour suppressor on chromosome 22. Suppose both *NF2* and this other tumour suppressor are mutated on one chromosome, and then LOH occurs, resulting in the other copies being lost. Both tumour suppressors would then be deactivated by only three hits. At least two such tumour suppressors are known, *SMARCB1* and *LZTR1* (see figure 1) [12, 13]. We will only consider *SMARCB1* here, and leave the inclusion of *LZTR1* to future work.

**Figure 1:**
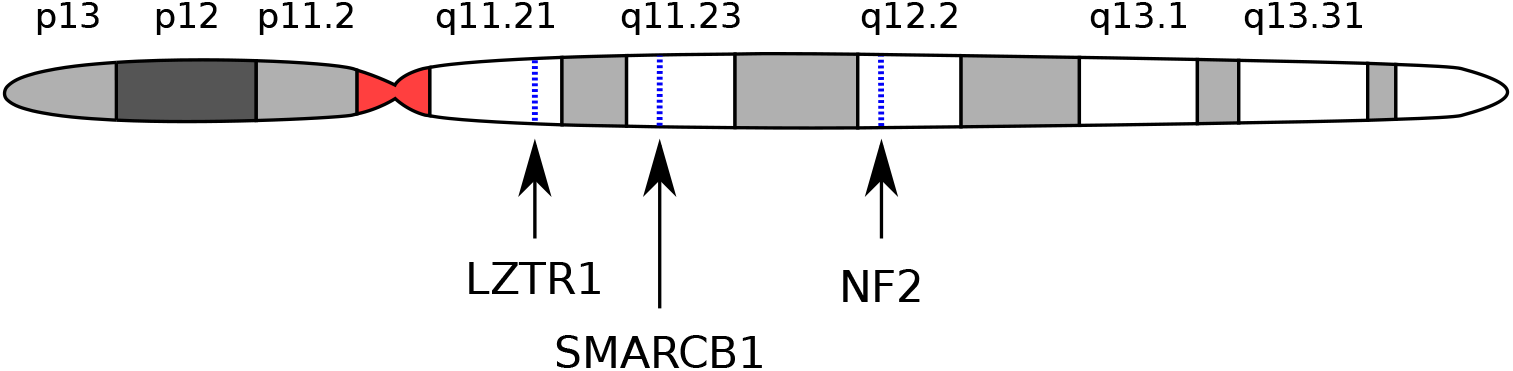
A map of chromosome 22 showing the approximate locations of *NF2* and two other tumour suppressor genes of possible interest, *SMARCB1* and *LZTR1* [12, 13]. With reference to UCSC Genome Browser [14].

We will first define our sporadic incidence model, before showing how to infer its parameters in section 2.1. Our goal will be to derive improved estimates for the underlying parameters from experimental data. These parameters will be critical to our models of malignancy in sections 3-4.

The model has four basic types of parameter: the base population *N*_0_ of progenitor cells from which the tumour derives; the mutation rates *μ_gene_* for point mutations on each gene, and the rate *r_LOH_* of LOH on chromosome 22; the selective advantages *s_i_* associated with mutant variants; and patient age *t* [10]. There is a lack of evidence for an exponential phase in the incidence of benign VS [4]. As a result, we will assume that the three initiating mutations are neutral (*s_i_* ≈ 0) [4, 11]. This reduces the parameters to the initial population *N*_0_, the relevant mutation rates *μ_gene_* and *r_LOH_*, and age t, the only independent variable [10].

The three alterations in this model consist of two hits to *NF2*, and one hit to either *SMARCB1* or a hypothetical oncogene. The model explicitly accounts for the different orders in which the hits can occur. There are three distinct sets of mutations that could result in a schwannoma, and each set may occur in many possible orders:

1. single copy inactivation of *NF2* → loss of 22q → gain of oncogene (3! = 6 different orders)
2. single copy inactivation of *NF2* → loss of second *NF2* allele → gain of oncogene (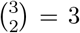 different orders)
3. single copy inactivation of *NF2* → loss of 22q → *SMARCB1* single copy inactivation (3! = 6 different orders)

In principle, a complete model should include hits to *LZTR1* as well as *SMARCB1*. Due to a lack of available data, we are leaving this to future work.

The end state of any of these pathways is a population of tumour cells. These neoplastic mutants are labelled 1, 2, and 3 in figure 2. These states correspond to subtypes of schwannoma with distinct alterations present: LOH will be found in states 1 and 3; pathogenic variants of *SMARCB1* will only be present in state 3; and *NF2* will be doubly mutated in state 2. These different orderings imply the existence of many distinct subpopulations of cells. These intermediate steps, between wild-type progenitor cells and neoplastic tumour cells, are labelled in figure 2.

**Figure 2:**
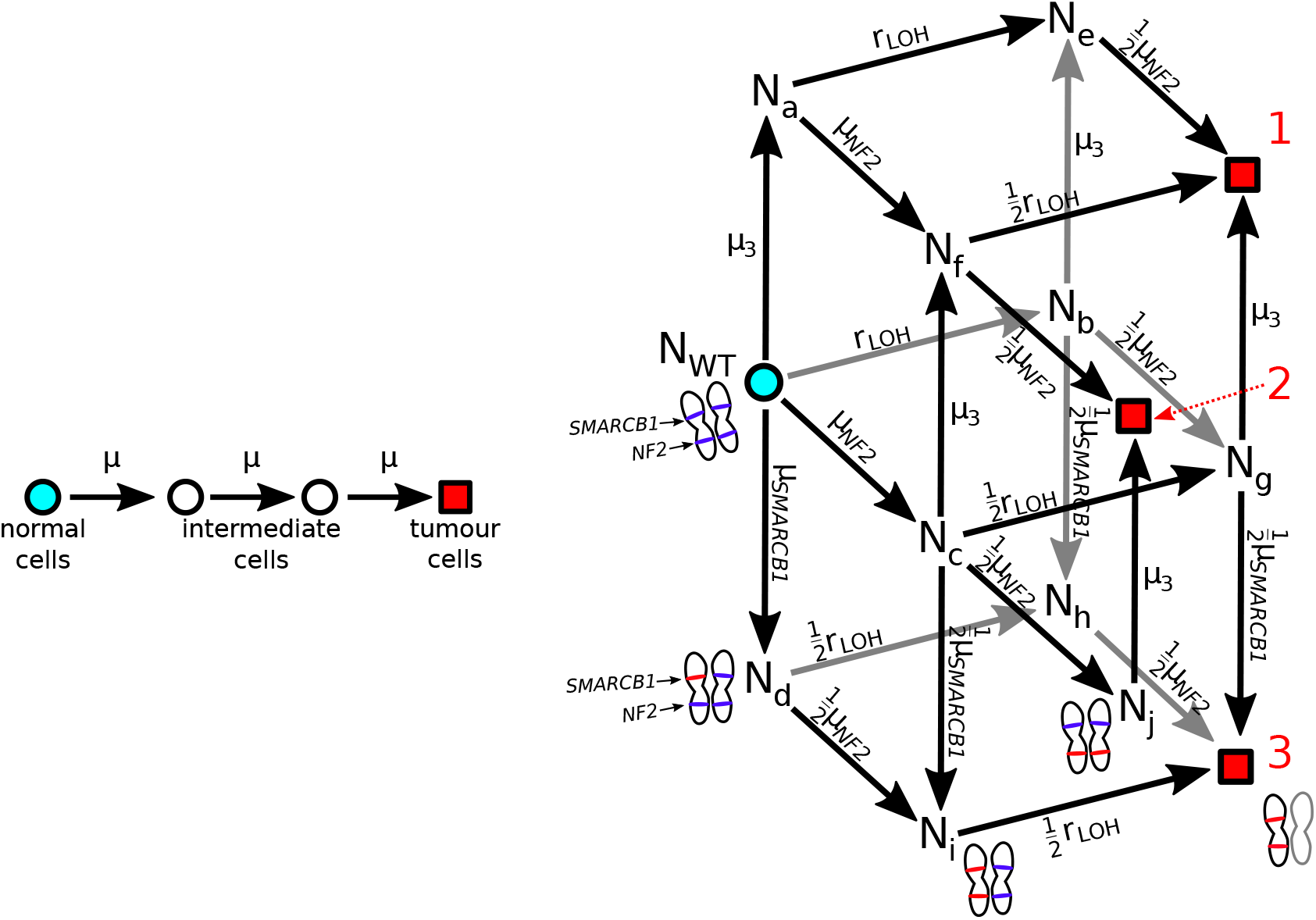
Left: a linear multi-stage model that does not attempt to distinguish between different kinds of alteration [4]. Right: the model in system (1) that allows for all possible orders of occurrence of the 3 alterations may be represented as a directed graph [10, 16]. All the Schwann precursor cells begin on the cyan circle, which represents the wild-type population *N_WT_*. The intermediate subpopulations are labelled with the corresponding *N_type_* from the system (1), and the mutation rates that connect them are labelled with *μ_gene_* and *r_LOH_*. The red squares 1, 2, and 3 represent neoplastic genotypes with different genetic alterations present.

**Figure 3:**
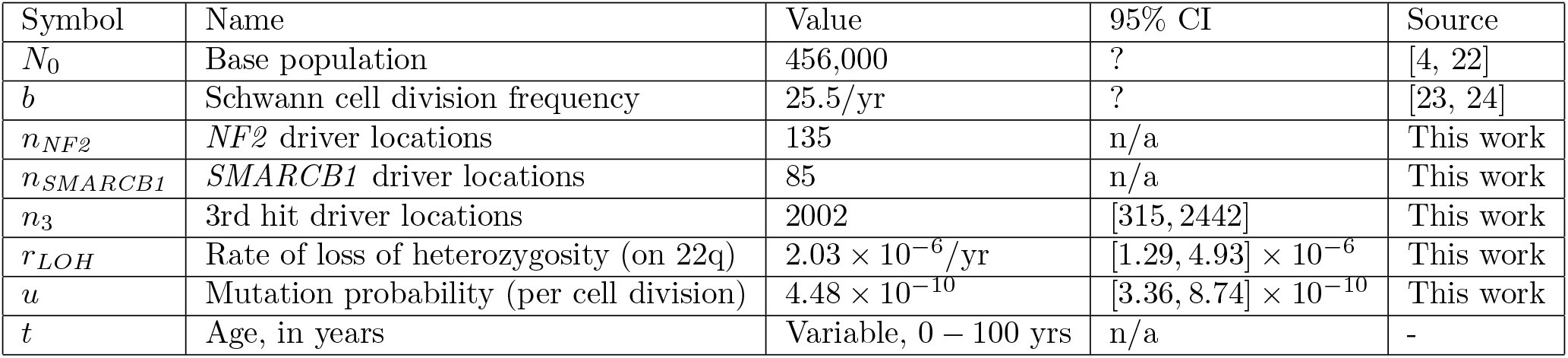
A summary of estimated parameter values for the sporadic vestibular schwannoma incidence model detailed in section 2.1. Uncertainties in the parameters *n_3_, r_LOH_* and *u* were derived by bootstrapping distributions from experimental results (section 2.2) [8, 12, 25].

At age zero, the entire population of *N*_0_ progenitor cells are assumed to start in the wild-type state, *N_WT_*. Random mutations result in intermediate mutants successively emerging, until a single neoplastic cell results. Rather than treat each subpopulation as a discrete random variable, we will employ a “mean field” or “fluid” approximation, and only track the *mean* subpopulation in each state [10, 11, 15].

Neoplastic cells emerge from the pre-neoplastic cells in a Poisson process. The rate of this process is the rate with which the relevant mutations appear in the subpopulations *N_e_* to *N_j_* (see figure 2, and system (2)). The subpopulations *N_m_* labelled in figure 2 follow a system of differential equations:

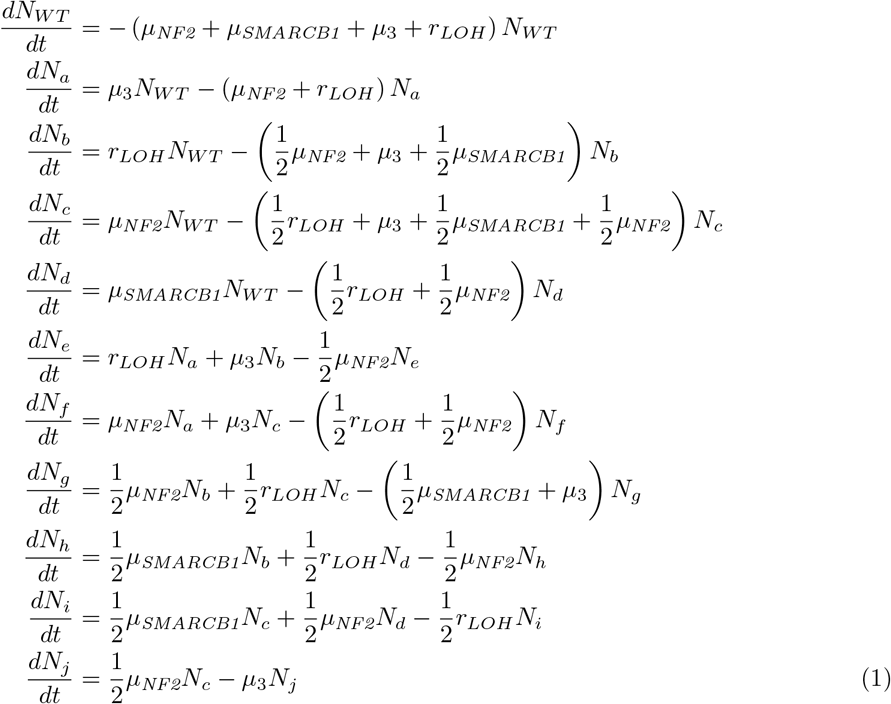

where the mutation rate *μ_gene_* for a given gene is determined by the error rate *u* per replication per base pair, the cell replication rate *b*, and the number of “sensitive locations” on that gene *n_gene_*, [10]:

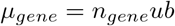

and *r_LOH_* is the rate of loss of heterozygosity on chromosome 22. In system (1), the subscript *μ_3_* stands for the mutation rate of hit 3, the hypothetical oncogene. The populations in system (1) have the initial conditions

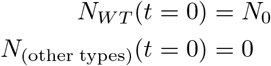

The probabilities of developing a schwannoma of subtype 1, 2 or 3 by age *t* follow

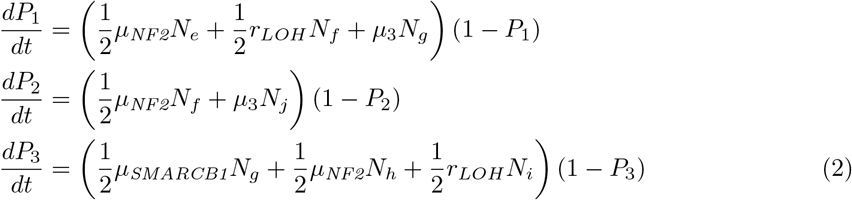

where *P_j_* is the probability for end-node *j* to be reached. These end nodes are labelled on figure 2.

Tumours in vestibular schwannoma often consist of multiple independent tumour foci on the same nerve, apparently polyclonal, that have collided to form a larger mass [17, 18]. This contrasts with models of tumour initiation in other diseases, where each tumour is assumed to be monoclonal [19, 20, 10]. For VS, the emergence of tumours of subtypes 1, 2, or 3 should therefore be treated as *independent* events, and not mutually exclusive events.

After solving systems (1) and (2) for *P*_1_, *P*_2_, and *P*_3_, we can use the independence of these events to find the probability to develop a schwannoma with LOH,

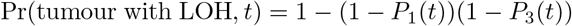

the probability to develop a schwannoma in which *SMARCB1* has been lost,

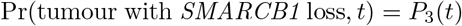

and the probability to develop a schwannoma with any of the above alterations,

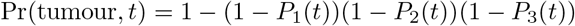

Vestibular schwannoma is a rare disease, so all the final probabilities *P_type_* must be small. When this is the case, we can make the approximation that quadratic and higher order terms (like *P*_1_ *P*_2_ or *P*_1_ *P*_2_ *P*_3_) are zero. The relevant formulae simplify considerably:

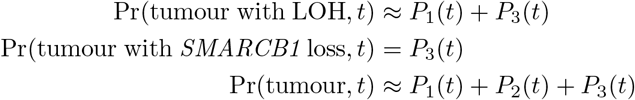

System (1) is linear in the intermediate populations *N_m_*, and can be readily solved by a variety of methods. Several underlying parameters are presently unknown, so we will first give a symbolic solution, and then determine numerical values of these parameters in section 2.1.

The intermediate populations of pre-neoplastic cells must also be small. When this is the case, each probability is approximately a power of age, *P_type_* ≈ *Ct^k^*. This can be seen from a power series solution of (1)-(2) around *t* = 0 [11, 10] (see appendix A). This is analogous to the classic “log-log” behaviour of many cancers [11, 21]. For sporadic vestibular schwannoma, this should be a good approximation at all ages (see appendix A for a detailed analysis). The solutions for *P*_1_, *P*_2_, and *P*_3_ read

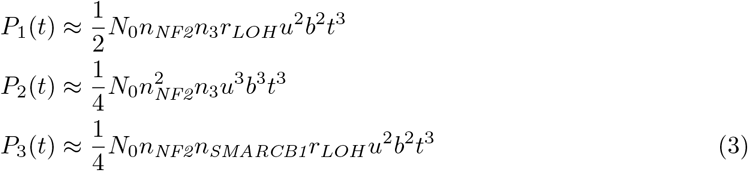

From (3), we can find the probability *P*(tumour, *t*) that a tumour has been initiated by age *t*:

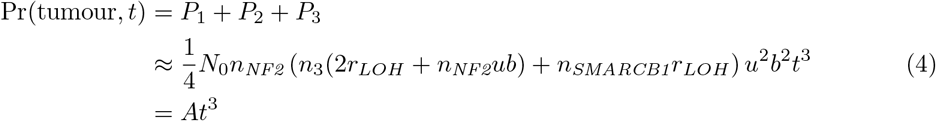

The constant *A* can be directly related to incidence data and other models, as shown in section 2.1 [4]. Also of interest are the probability to develop a tumour in which *SMARCB1* has been lost,

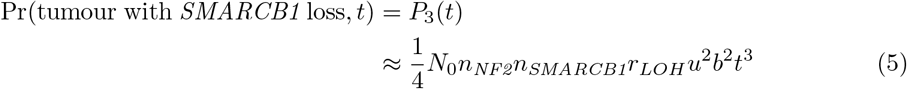

and the probability to develop a tumour that displays LOH on chromosome 22,

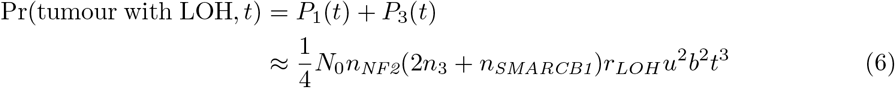

Equations (5) and (6) are crucial in determining the model parameters in section 2.1, as they can be related to empirical frequencies of pathogenic variants.

### 2.1 Parameter estimation

The model in section 2 has seven parameters: the initial wild-type population *N*_0_; the cell turnover rate *b*; the mutation rate per base pair *u*; the sensitivity parameters *n_NF2_, n_SMARCB1_*, and *n*_3_; and the rate of LOH *r_LOH_*. These must now be estimated.

#### 2.1.1 Precursor cell population and division rate

The base population of precursor cells *N*_0_ can be estimated from anatomical considerations. There are 2 vestibular nerves, with around 19,000 axons in each [4, 22]. The region on the vestibular nerve from which schwannomas usually originate is approximately 6mm long [4], and the approximate spacing between Schwann cells is approximately 0.5mm [4]. Thus, the base population in the normal tissue pool *N*_0_ is approximately

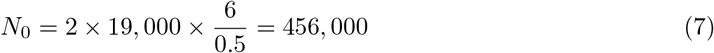

This is higher than R. Woods’ 2003 estimate [4], which we attribute to having a more recent estimate for the number of axons in each vestibular nerve [22].

The rate of cell division *b* in the precursor cell population can be estimated from an experiment on Schwann cell proliferation following injury [23]. Their observed rate of Schwann cell proliferation of 7%/day corresponds to a value of *b* of 25.5/year.

#### 2.1.2 Sensitive sites

The effective numbers of sensitive locations *n*_*NF*2_ and *n_SMARCB1_* can be calculated from reference sequences [10, 26, 27]. A site is “sensitive” if a mutation there results in a nonsense codon, truncating the gene (see figure 4). Only sites on the longest open reading frame are considered. Express *n_gene_* as

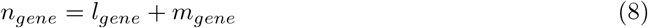

where *n_gene_* is the overall number of sensitive sites, *l_gene_* is the number of substitution-sensitive sites, and *m_gene_* is the number of indel-sensitive sites.

To calculate the number of substitution-sensitive sites *l_gene_*, count the number of places where a base substitution could result in a nonsense codon. If two possible substitutions can result in a stop codon, count this base twice. Finally, multiply this by 1/3 to account for the fact that only one of the three possible substitutions will result in a nonsense codon.

To calculate *m_gene_*, the empirical approximation

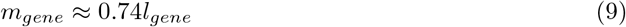

from Bozic et al. 2010 can be used [28].

**Figure 4:**
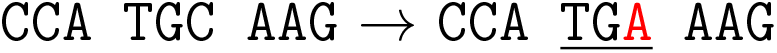
A diagram of a truncation mutation: substitution of a single base pair can result in a premature stop codon.

**Figure 5:**
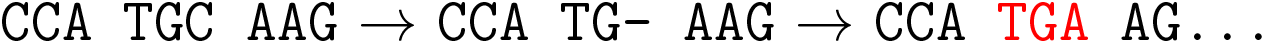
A diagram showing how a small deletion of a single base pair (length *k* =1) can result in a premature stop codon. The deleted site is “sensitive”, and will result in a loss of function.

A more mechanistic way of calculating *m_gene_* is desirable, as this will be an important parameter in section 4, so we will outline a similar counting argument. As a first approximation, we will *only* count sites as sensitive if a deletion there is next to or overlaps a nonsense codon (see figure 5) and assume this *always* causes a loss of function. We can now count the number of sites *m*(*k*) where a deletion of length k results in a stop codon: `TAG, TAA`, or `TGA`. Small insertions and deletions have a distribution of possible lengths *k*, with *k* = 1 occurring more often than any other length [29, 30, 31].

Calling the length of an indel *k*, with *k* > 0 corresponding to deletions, and *k* < 0 to insertions; denote its frequency of occurrence *f*(*k*). For each possible length *k*, there is a corresponding number of sensitive sites *m*(*k*) on the gene. This is calculated in a similar way for each *k*: for each site on the gene, check if deleting the base pair there, and *k* – 1 bases afterwards, results in a nonsense codon. The number of indel-sensitive sites, *m_gene_*, is then the average of *m*(*k*) over the length distribution *f*(*k*):

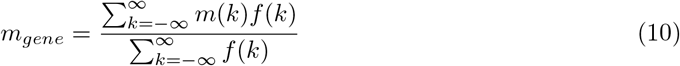

In the following, we will assume that insertions are just as likely as deletions, and *f*(*k*) is symmetric. Given a sequence, we can calculate *m*(*k*) explicitly for all values of *k*. Then, given a length distribution *f*(*k*) such as

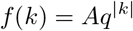

*m_gene_* can be computed explicitly given a sequence. We will use *q* = 0.53 for consistency with [29], Supplementary Table 5A. Our results were largely insensitive to the exact choice of *q*. Supplemental Python code that computes *l_gene_* and *m_gene_* from EMBL sequences has been provided.

Using reference sequences for *NF2* and *SMARCB1* from the European Nucleotide Archive [26, 27], we find:

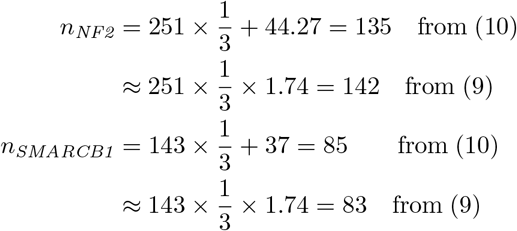

#### 2.1.3 Base pair mutation rates, rates of LOH, and *n*_3_

The number of sensitive sites *n*_3_ on the oncogene (or proto-oncogenes) cannot be estimated by a similar counting argument, because the identity of this gene is unknown. Direct measurements of the rate *r_LOH_* of loss of heterozygosity in Schwann cells, and of the per-base mutation rate *u*, are also unavailable.

However, all three parameters can be constrained using measurements of the empirical probabilities *f_LOH_* of LOH, and *f_SMARCB1_* of pathogenic variants of *SMARCB1*. These can be computed from experimental studies [25, 12, 32]. In a sample size of n patients, if *k_LOH_* patients are found to have LOH on 22q, then the relative frequency *f_LOH_* can be estimated:

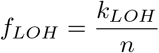

i.e. the marginal total of patients with LOH divided by the total number of patients [32]. A similar calculation can be done for *f_SMARCB1_*, the frequency of pathogenic variants of *SMARCB1*.

However, there is a flaw with this naive approach: there is a chance that any experiment with a finite sample size n will contain no positive samples, and *k_LOH_* or *k_SMARCB1_* may be zero purely due to stochastic sampling error. We shouldn’t conclude as a result that the empirical probability *f* is actually zero: there is in fact a range of credible values consistent with *k* = 0 to a given degree of confidence [33, 34, 35]. The same would be true if LOH was found in every sample, and *k_LOH_* = n. To address this problem, we use additive smoothing with a pseudocount of 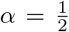, corresponding to the Jeffreys prior [36, 34]:

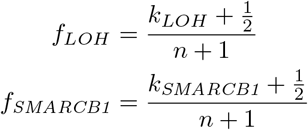

Further discussion of this choice can be found in appendix B.

Both *f_LOH_* and *f_SMARCB1_* can be predicted by our model. From (4), (5), and (6), theoretical predictions for *f_LOH_* and *f_SMARCB1_* can be found:

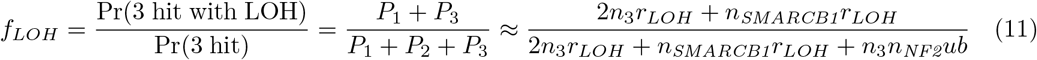

and

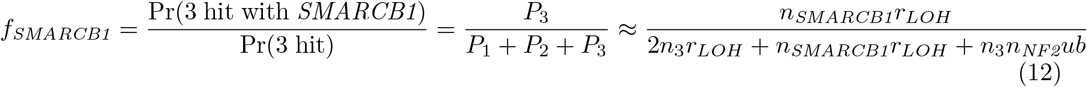

*f_LOH_* and *f_SMARCB1_* are algebraically independent. So, if *f_LOH_* and *f_SMARCB1_* are known experimentally, then two parameters of the model can be fixed. *n*_3_ and *r_LOH_* can be expressed explicitly in terms of *f_LOH_* and *f_SMARCB1_*:

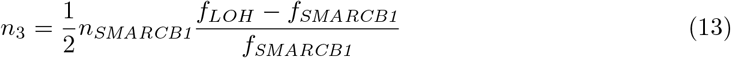

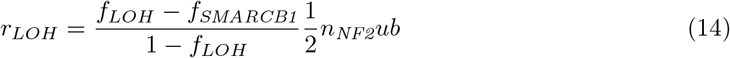

so that if *f_LOH_* and *f_SMARCB1_* are both known, then the only remaining free parameter is *u*.

A recent study of *NF2* status and LOH in sporadic vestibular schwannoma, which also tested for copy-neutral LOH, found that out of 23 patients, 17 had some form of LOH [25]. So, *f_LOH_* = (17 + 0.5)/(23 + 1) = 73%.

Relevant measurements of *SMARCB1* status were not available, so we performed our own sequencing experiments to determine *f_SMARCB1_* Sequences from 32 sporadic vestibular schwannomas were sourced from the NHS DNA archive in the Manchester Centre for Genomic Medicine, and pathogenic variants of *SMARCB1* were searched for using Sanger sequencing.

Sanger sequencing of *SMARCB1* involved initial amplification of each coding exon, including 50-100bp flanking intronic sequence per exon, using GoTaq G2 (Promega, Madison, WI, USA). The products were purified using Ampure cleanup beads and the purified products were used as a template for sequencing PCRs using BigDye Terminator v3.1 (Life Technologies, Paisley, UK). Sequencing PCR products were analysed on an ABI3730xl DNA Analyser (Life Technologies, Paisley, UK). Sequencing chromatograms were aligned to the reference sequence to identify variants.

Our experimental results found no pathogenic variants of *SMARCB1* in a sample of 32 sporadic vestibular schwannomas. The naive estimate of *f_SMARCB1_* is therefore *k/n* = 0/32 = zero. This presents a problem, as our estimate for *n*_3_ diverges when this is the case. As discussed above, we use additive smoothing to regularise *f_LOH_*, with a pseudocount of 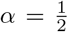 (see appendix B for a further discussion) [34]. Thus,

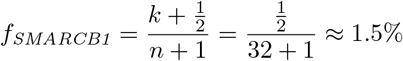

or *f_SMARCB1_* = 1.5%.

The same quantity *f_SMARCB1_* has also been measured for both sporadic schwannomatosis, a genetic predisposition to schwannoma that involves *SMARCB1* [12]; and for sporadic *spinal* schwannomas [13]. It is instructive to compare *f_SMARCB1_* values across the three diseases.

K. Hadfield et al. found that of 28 patients with schwannomatosis, 2 had mutations on *SMARCB1*, so *f_SMARCB1_* = 2.5/29 = 8.6% [12]. Comparison of spinal and vestibular schwannoma shows broadly similar frequencies of both LOH and *SMARCB1* loss across the three different types, generally less than 10% (see figure 6) [25, 12, 8].

**Figure 6:**
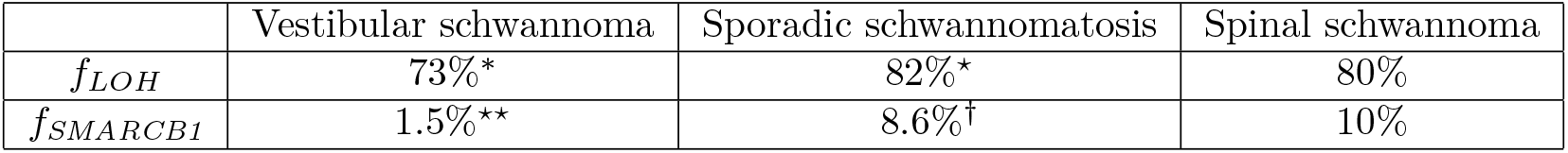
Estimates of the frequencies *f_LOH_* of LOH and *f_SMARCB1_* of pathogenic mutant *SMARCB1* alleles in different studies. *: from M. Carlson et al. 2018 [25]; ⋆: from K. Hadfield et al. 2010 [8]; †: from K. Hadfield et al. 2008 [12]; ⋆⋆: this work. Both spinal schwannoma estimates are from I. Paganini et al. 2018 [13]. All the *f_SMARCB1_* estimates are in the range 0 – 10%. It may be noted that by the “rule of three” [35] that the 95% confidence interval on our *f_SMARCB1_* estimate is [0, 9.4%], suggesting that the differences in *f_SMARCB1_* between the diseases may not be statistically significant.

Together with *n_SMARCB1_* = 85, the values of *f_LOH_* = 73% and *f_SMARCB1_* = 1.5% imply that

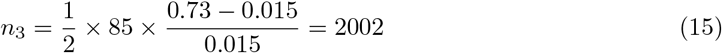

from (13), and

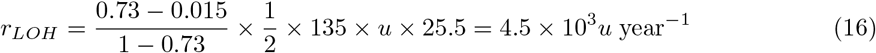

from (14). The only remaining degree of freedom is now u, the error rate per base pair per division. We determine *u* first by fitting a power law,

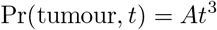

to incidence data from D.G.R. Evans 2005 [1]. The fitting constant *A* is related to the model parameters by equation (4),

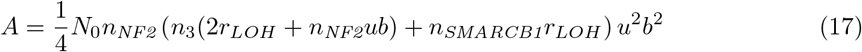

The probability to develop a tumour by age *t, P*(tumour, *t*), is the cumulative incidence by age *t*, after correcting for survival. Before fitting, the empirical incidence was therefore corrected for mortality using life tables for the period and region of the study [37].

The dataset was also truncated above the age of 80. This is because the relevant population size at ages greater than 80 was very small, and the incidence displays an old-age “plateau” that we suspect is an observational artifact (DGR Evans, private correspondence, 2019). This amounts to excluding 8 patients from the original dataset of 417, so only 2% of the data has been trimmed [1]. In addition to these modifications to the dataset, our model only accounts for VS associated with somatic *NF2* loss. This occurs in 85% of cases, so the cumulative incidence data is also rescaled by 85% [3].

The resulting best-fit value was *A* = 2.26 × 10^−11^ /yr^3^, with a standard error of *σ*(*A*) = ±2.95 × 10^−13^. This was not an important source of uncertainty, and was dwarfed by the uncertainty in *n*_3_ and *r_LOH_* due to the small sample sizes of the experiments used to determine alteration frequencies (see section 2.2).

This three-hit model displays a high goodness of fit, with *R*^2^ = 0.989. However, prior work had already established that a three-hit model provided a good fit to this dataset, so replicating this finding with high confidence is not statistically remarkable [4]. The purpose of this procedure was primarily to provide new parameter estimates. These new estimates, in particular *u* and *r_LOH_*, will be useful in our study of malignant transformation (section 3).

Together with equation (17), this value for A implies that

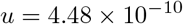

per base pair per division. This is similar in order of magnitude to the value of *u* ≈ 10^−9^ estimated by C. Tomasetti and I. Bozic [38]. This estimate for u now allows a concrete numerical value to be placed on *r_LOH_*:

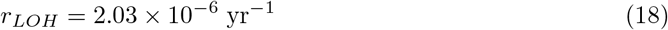

This is rather low in comparison to Paterson, Bozic and Clevers’ estimate for *r_LOH_* = 1.2 × 10 ^−4^/yr in colorectal cancer [10]. Possible explanations for this difference are discussed in section 5.1.

### 2.2 Uncertainties in parameters

The sample sizes used to estimate frequencies of *f_LOH_* and *f_SMARCB1_* were relatively small, both with *N* = 20 to 30 patients. As a result, *n*_3_, *u*, and *r_LOH_* have important sources of uncertainty that are difficult to quantify with standard methods – i.e. propagation of uncertainty typically assumes normally distributed errors, which is not a valid assumption when sample sizes are small [40].

To address this, distributions for these parameter values were bootstrapped. Randomly resampling regularised datasets for LOH and *SMARCB1* with replacement generates a distribution of estimated values for *n*_3_, *u*, and *r_LOH_* (see appendix B) [25, 12, 41]. A 95% confidence interval can then be determined for each parameter from the resulting distribution [39]. These confidence intervals read:

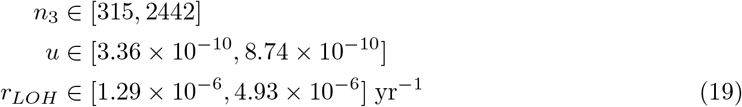

In the context of bootstrapped distributions, the “best estimates” from section 2.1 can be interpreted as modal estimates. Supplemental Python code that implements the bootstrapping procedure has been provided. All parameter estimates and intervals are presented in table 3 and figure 7.

**Figure 7:**
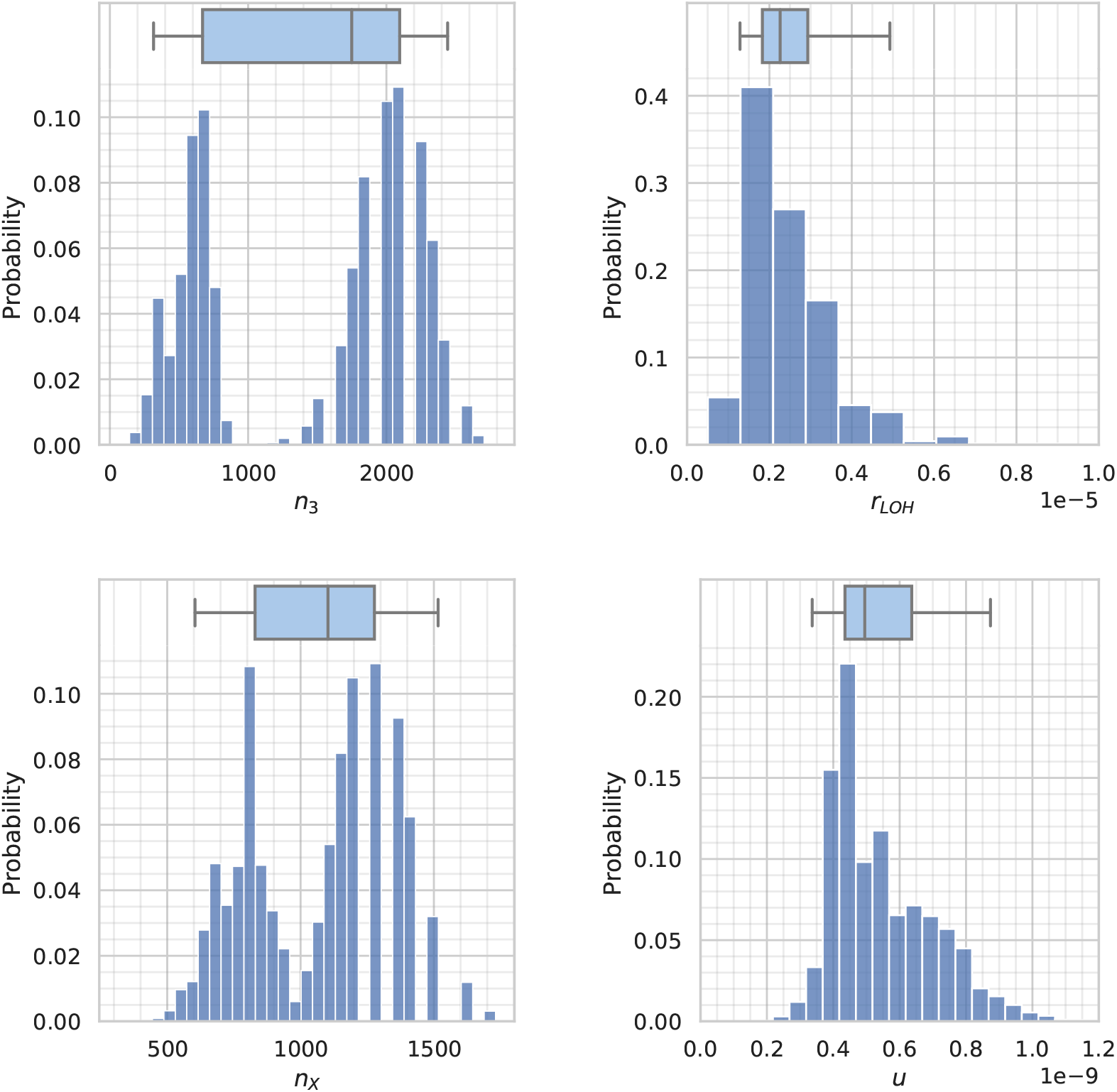
Bootstrapped distributions for the three variables *n*_3_, *r_LOH_*, *n_X_* (see section 3.2), and *u*. These were determined by the frequency of LOH from [8], by our own attempted measurements of pathological *SMARCB1* variants, and by incidence data from [1]. Equations (13), (14) and (17) allowed estimates of *n*_3_, *r_LOH_* and u to be generated. The box represents the median estimate and inter-quartile range; the outer whiskers represent the 2.5% and 97.5% quantiles, i.e. the 95% confidence interval [39]. Note that the distributions for *n*_3_ and *n_X_* appear to be bimodal.

## 3 Modelling sporadic malignant transformation in schwannoma

Additional mutations may occur as the tumour expands. When the “right” mutations occur together, a cell within the tumour gains a malignant phenotype [42]. In the case of malignant schwannoma, it is not currently clear whether these additional hits are due to oncogenes, tumour suppressors, or a combination of the two. We investigated several possibilities.

It was found during preliminary work that if malignant schwannoma were caused by a single oncogene, then the chances of its emerging from any benign schwannoma would be extremely high, on the order of 100%. A similar conclusion was also reached for any tumour suppressor on chromosome 22. As malignant schwannoma is a rare disease, it is far more likely that these hits involve multiple hits – the simplest hypothesis is that a tumour suppressor gene that is *not* on chromosome 22 is responsible.

In the following, we call this hypothetical tumour suppressor gene “ *TSX*”. We should bear in mind that there may be *several* such tumour suppressors, presumably associated with distinct molecular subtypes of malignant schwannoma. But for simplicity, we will develop the simplest model that is consistent with experimental observations, and make the operational assumption that there is only one such gene. Models that fully account for genetic diversity and multiple tumour suppressors will be left to future work. If the gene *TSX* needs to be extremely large for the model to make realistic predictions, this would be an early warning sign that we have missed some of this genetic diversity.

Once again, we will adopt a mean-field approach, and model the mean of intermediate populations [10]. Unlike in our approach in section 2, the underlying “wild-type” population is not constant, but expanding as the tumour grows [11, 43].

We will make the approximation that there is no cell death in the tumour, and all growth occurs at the outermost edge [43]. This is likely to be an underestimate of the real rate of cell division and turnover. But it allows us to eliminate dependence on the time *t* elapsed since tumour initiation, which is not directly observable. This obviates any assumptions about the tumour’s growth curve.

In each cell division event, there is some probability to create a mutant daughter cell. This can occur through either a point mutation on *TSX* with probability *n_X_u*, or through LOH on the relevant chromosome with probability *p_LOH_*. The subpopulations *N_k_*, *N_l_, N_m_* in the expanding tumour (see figure 8) obey the following differential equations:

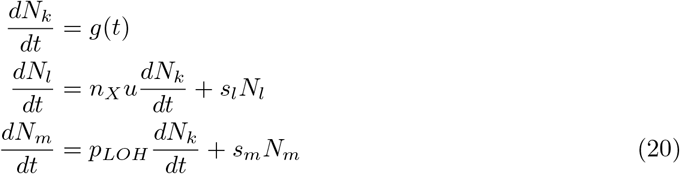

where the growth rate *g*(*t*) of the tumour is some smooth function of time with *g*(0) = 0, and *s_l_* and *s_m_* are fitnesses of the mutant clones. System (20) has the initial conditions:

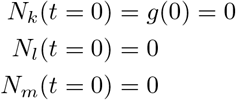

so that *N_k_* is just

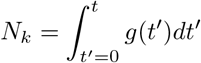

**Figure 8:**
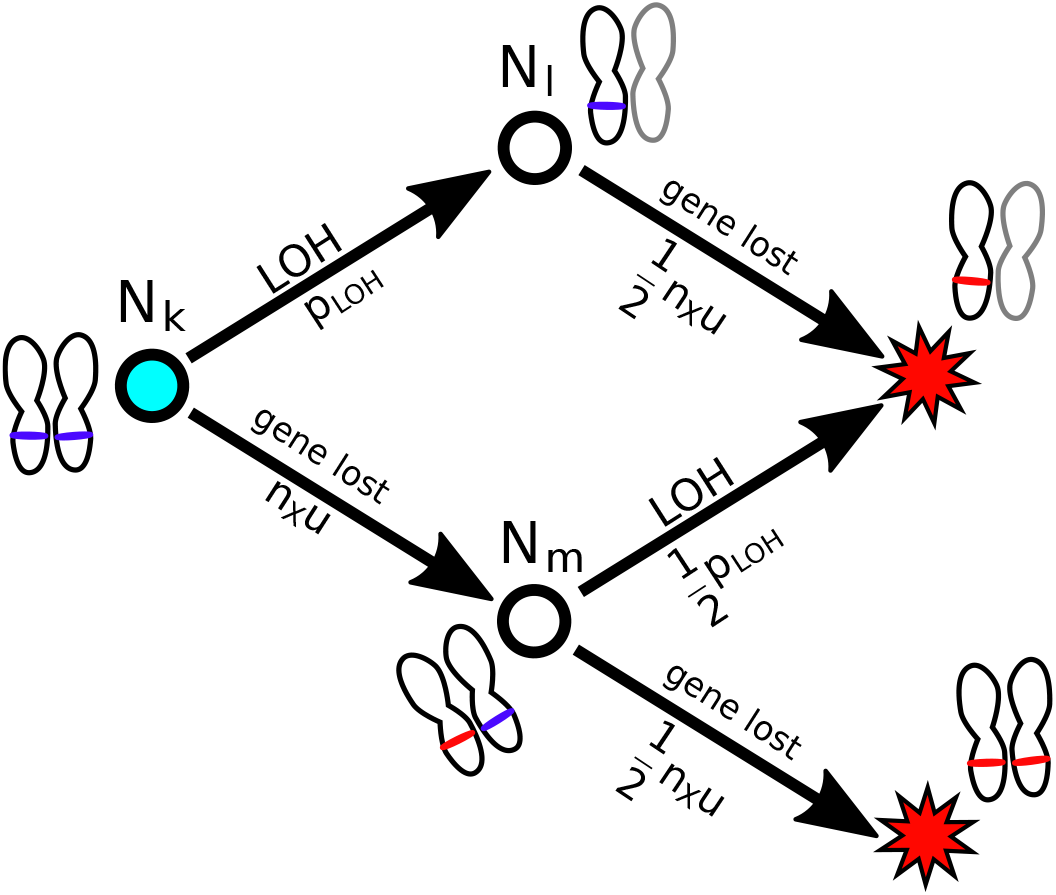
Our 2-hit model of tumour suppressor inactivation, accounting for all three orders of appearance. The cyan circle is the genotype of the initial, benign tumour. The red sites represent the malignant genotype. The subpopulations from system (20) are labelled *N_k_, N_l_, N_m_*, and the mutation probabilities indicated below the corresponding arrows.

**Figure 9:**
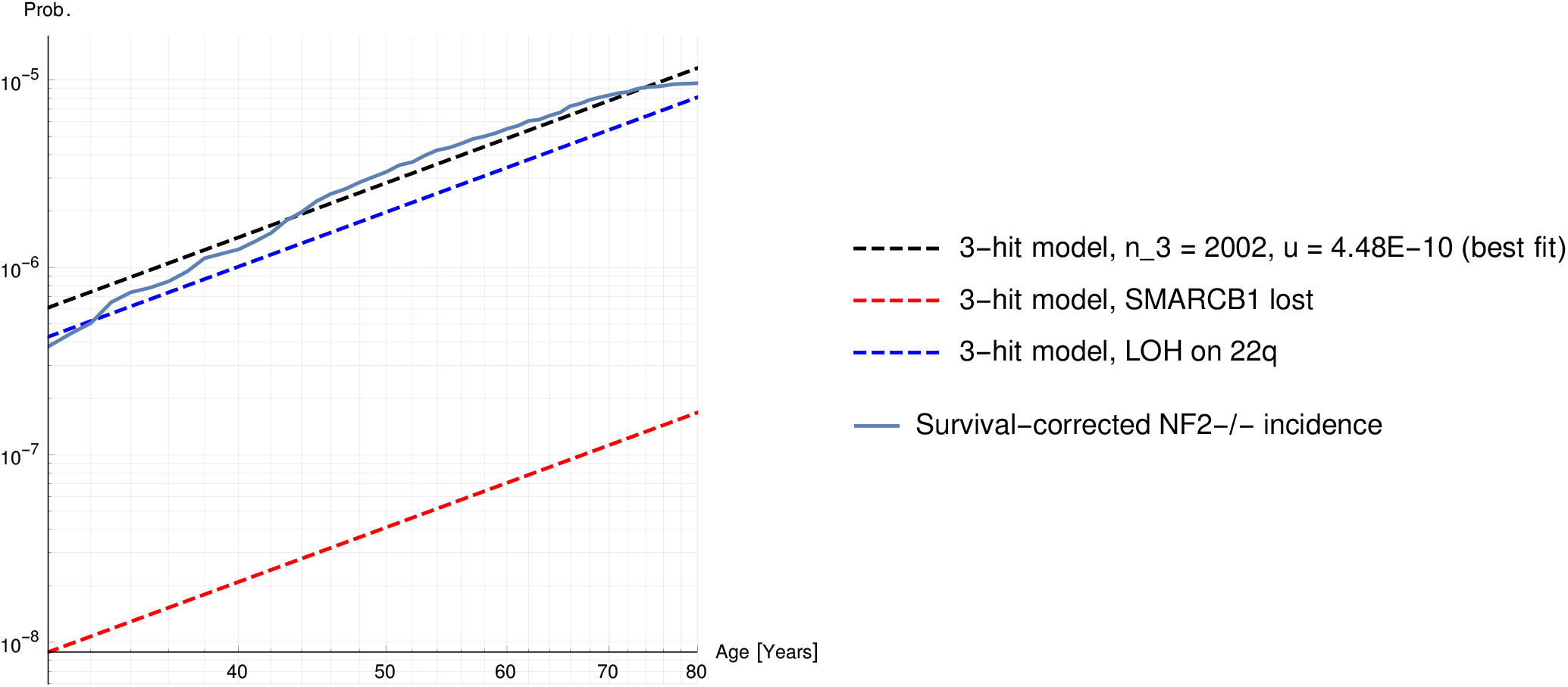
The probability of developing sporadic vestibular Schwannoma (dashed curves) and empirical cumulative incidence (solid line) as a function of age. The multi-stage model from equation (4) is shown with a black, dashed curve, and has parameters *u* = 4.48 × 10^−10^, the number of sensitive sites on the third hit *n*_3_ = 2002, and *r_LOH_* = 2.03 × 10^−6^yr^−1^. The blue dashed curve shows the modelled risk of developing a Schwannoma with LOH on 22q; and the red dashed curve shows the risk of developing a Schwannoma with a pathogenic variant of SMARCB1. Confidence intervals for the model parameters are established in section 2.2 and given in table 3. The empirical curves were derived from incidence data in [1] and corrected for mortality using life tables for the same period from [37].

Note that the populations are easier to measure than tumour age *t*.

As in the model from section 2, we will assume that as soon as a malignant cell emerges in the benign tumour, it immediately establishes a growing malignant lineage. The probabilities for a malignant clone to emerge with and without LOH follow

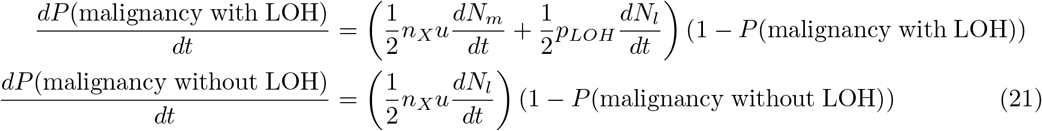

respectively, where *p_LOH_* is the probability for LOH to occur during a single division, and *n_X_* is the number of sensitive locations on *TSX*. For the parameters *u* and *p_LOH_*, we will use the value of *u* = 4.48 × 10^−10^ from section 2; for *p_LOH_*, we will use a value consistent with our previous *r_LOH_* estimate:

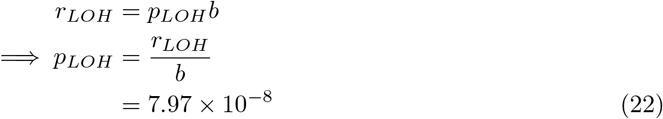

In the absence of better experimental constraints, we will take *p_LOH_* to be the same for all chromosomes. As *TSX* is currently unidentified, *n_X_* cannot be determined *a priori* from a reference sequence like in section 2.1. We constrain *n_X_* in section 3.1.

General solutions to system (20) can be written:

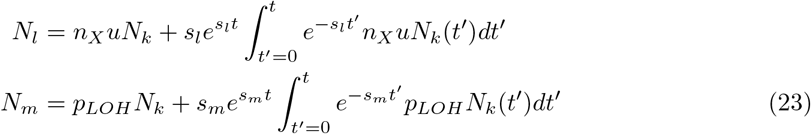

The fitnesses *s_l_* and *s_m_* relative to the “wild-type” tumour cells *N_k_* are not known. Since both subpopulations *l* and *m* still have one active copy of *TSX*, we will assume these mutations are neutral, *s_l_* = *s_m_* = 0. This amounts to assuming that *TSX* is haplosufficient. In this limit, (23) becomes

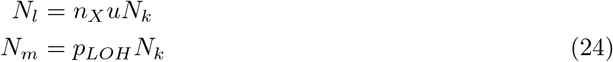

Note that *all* explicit dependence on t has disappeared, and these depend purely on *N_k_*. The risks of different types of malignancy are then given by the solutions of (21),

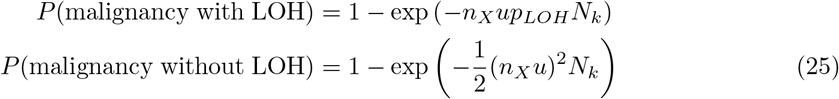

The overall risk of malignancy can be calculated from (25) using the relation

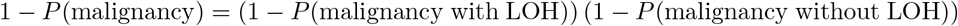

from which it follows that

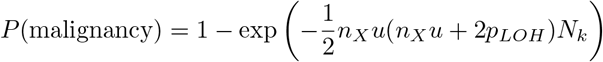

We can also note that both *n_X_u* and *p_LOH_* are small, on the order of 10^−7^. The tumour population N and subpopulation *N_k_* must therefore be similar, to within one part in 10^−7^:

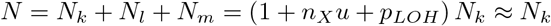

so as a result,

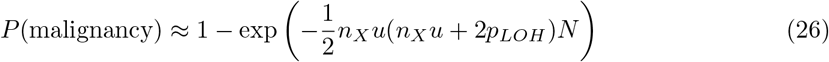

Finally, we can notice once again that because malignant Schwannoma is rare [2], *P*(malignancy) must be very small, *P* ≈ 0.2% ≪ 1, and (26) simplifies to

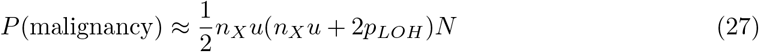

To connect *P*(malignancy) to the observed volume *V* of the tumour, the population *N* must be related to *V*:

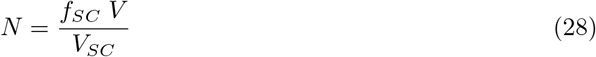

where *f_SC_* is the fraction of cells in the tumour that are Schwann cells, *V_SC_* is the volume of an individual Schwann cell (in mm^3^), and V the volume of the tumour (also in mm^3^). The majority of cells in schwannomas are not Schwann cells, but macrophages: the best available estimates of *f_SC_* are in the range 0.3-0.5 [44]. Of these, we choose the upper value of *f_SC_* = 0.5. A representative value for the volume *V_SC_* can be set at 1.6 × 10^−6^ mm^3^ (from [45]).

The final expression for the risk of malignancy in terms of tumour volume is thus

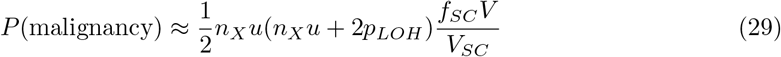

The tumour volume *V* can be observed, in principle, on MRI scans [46]. Notably, the risk of malignancy is simply proportional to tumour volume *V*. We constrain the number of sensitive sites *n_X_* in section 3.1, and explore a range of plausible values in figure 11.

**Figure 10:**
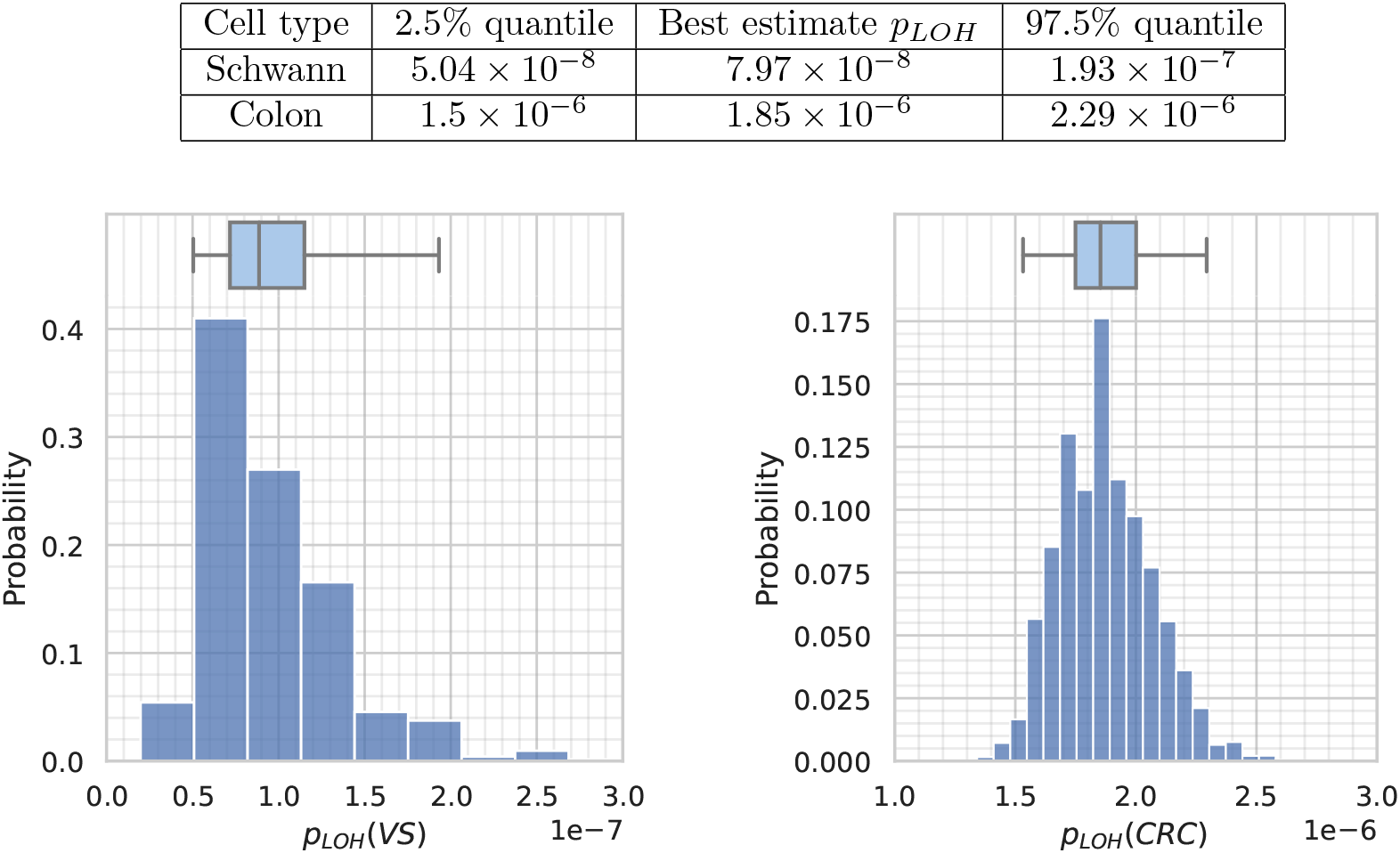
Quantiles for bootstrapped distributions for *p_LOH_* estimates (above), and visualisations of the distributions for Schwann progenitor cells (below left), and colorectal crypt progenitor cells (below right), demonstrating a substantial difference in order of magnitude. The estimates for colon cells were derived from Huang et al. 1996 [68] and Paterson et al. 2020 [10]. The estimates for Schwann cells are original to this work.

**Figure 11:**
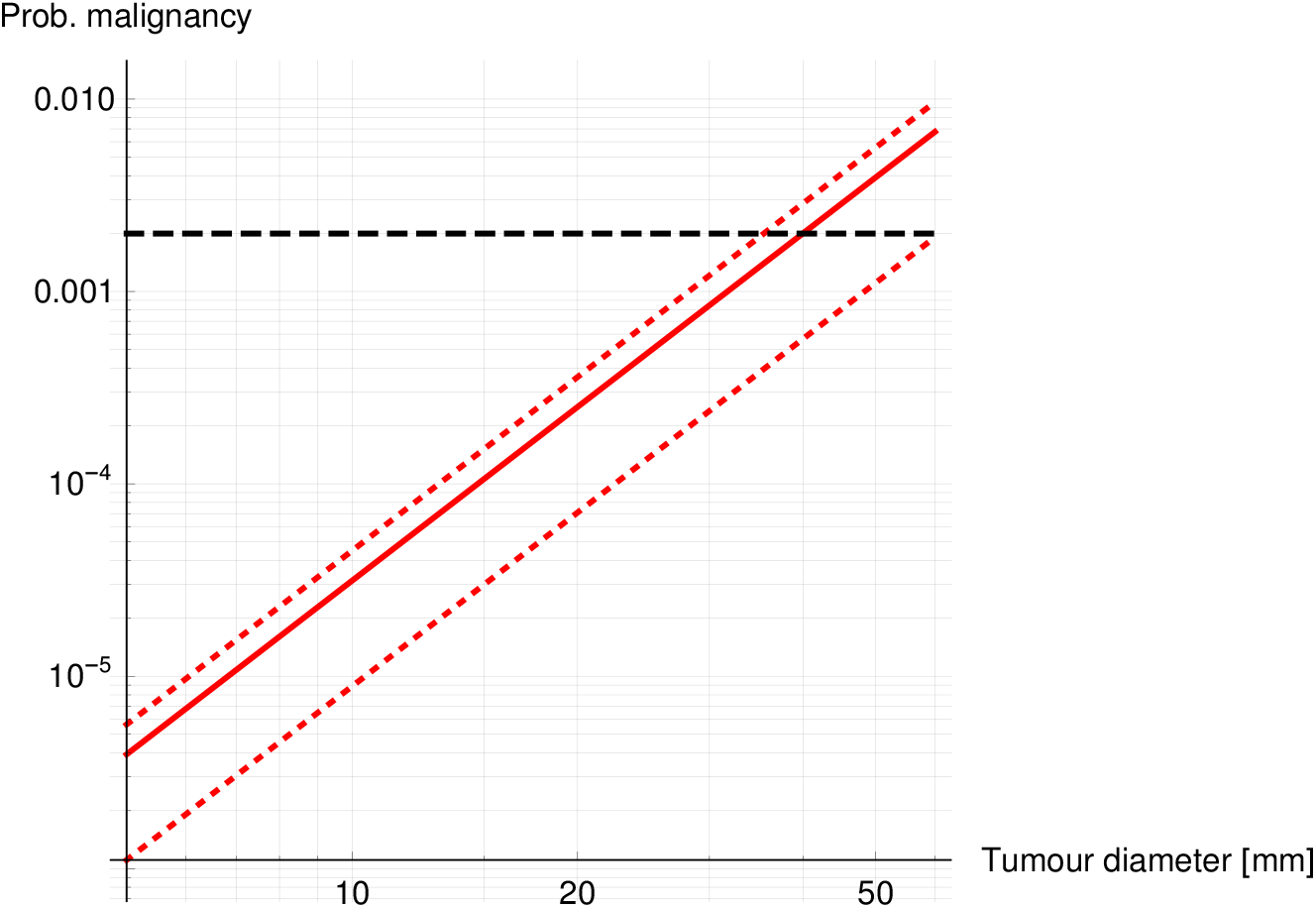
Risk of spontaneous malignant transformation in a benign schwannoma as a function of diameter *d* in mm, as modelled by equation (29). The estimated lifetime risk ≈ 0.2% is again shown as a horizontal dashed line [2]. The risk curve for our best estimate for *n_X_* = 1245 is shown with a solid red line: the upper and lower confidence intervals of 1515 and 604 with dotted red lines, above and below (from section 3.1)

### 3.1 Parameter estimation

The main unknown in this model of malignant transformation is *n_X_*, the number of sensitive sites on *TSX*. As the identity of this gene is unknown, *n_X_* cannot be calculated using a reference sequence. An estimate could help to constrain its identity.

From the SEER study, we know that the lifetime risk of malignant schwannoma ≈ 0.2% [2]. The model in section 3 avoided using “tumour age” as an independent variable, which is usually unobservable: the main result, equation (29), is formulated in terms of tumour volume.

If a typical tumour diameter on surgery is 40mm, the tumour will contain roughly *N* ≈ 10^10^ cells [47, 48]. If *P*(malignancy) ≈ 0.2% [2], then

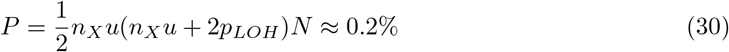

from equation (26). Rearranging (30) for *n_X_* is elementary. Given that *u* = 4.48 × 10 ^−10^, *N* ≈ 10^10^, and *p_LOH_* = 7.97 × 10^−8^, *n_X_* should then be

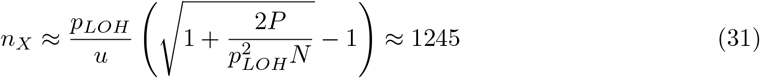

While this is a high figure, it is not impossible for *TSX* to be one gene: one may compare *n_APC_* = 604 from [10]. It nonetheless seems more likely that several tumour suppressors are involved, and this high estimate for *n_X_* indicates a degree of genetic diversity.

This estimate must be qualified by the uncertainties in *u* and *r_LOH_*. We can place confidence intervals on *n_X_* by a similar bootstrapping procedure to that in section 2.2. Using equation (31), one *n_X_* estimate can be determined for every pair of *u* and *r_LOH_* values. A distribution for *n_X_* can therefore be generated at the same time as those for *u* and *r_LOH_*. The resulting 95% confidence interval is

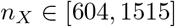

The full bootstrapped distribution for *n_X_* is shown in figure 7. Supplemental Python code has also been provided.

### 3.2 LOH in malignant schwannoma

Another testable prediction is the proportion of malignant tumours that should display LOH. Considered as a subtype of schwannoma associated with a mutation on *TSX*, levels of LOH on chromosome 22 should be similar to that of benign tumours. If malignant transformation involves an additional tumour suppressor gene, and it is not on chromosome 22, we should also expect to see LOH elsewhere.

This “excess” LOH should be found on the chromosome that carries *TSX*, the hypothetical tumour suppressor from section 3. Studying LOH, especially in the form of copy number changes, has already been widely used to search for tumour suppressors, although it only gives limited information about the precise locus [49]. In this section, we estimate the frequency *f_LOH_* of LOH aggregated across all chromosomes, relate this to *n_X_*, and estimate a minimum useful sample size for a study.

The parameter *n_X_* can be mathematically related to *f_LOH_* by a similar method to that used for *n*_3_ in section 2.1. This will be useful for two reasons. Firstly, so that future measurements of *f_LOH_* can be used to estimate *n_X_* [49]. Secondly, *f_LOH_* can be estimated in advance to judge whether or not such an experiment is worthwhile.

From equations (24) and (26), it follows that

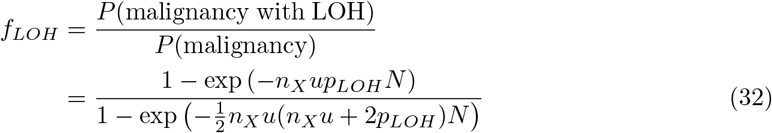

where *f_LOH_* is the excess LOH found at the locus of the unknown tumour suppressor. Again, we know that malignancy is very rare, with a lifetime risk of around 0.2% [2]. So once again, *P*(malignancy) must be very small. In this limit, the dependence on *N* drops out, and equation (32) simplifies to

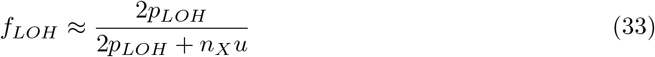

Although we knew that *P* was small empirically, strictly speaking, (33) can only be a good approximation of (32) when *N* is sufficiently small. Substituting our estimates of *u*, *p_LOH_* and *n_X_* into (27), one can show that this holds when

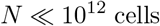

A tumour with 10^12^ cells would correspond to a tumour 18cm in diameter. This is 100 times larger than the majority of either malignant or benign schwannomas [48, 47]: it should not even be possible for a tumour this large to form inside the cerebellopontine angle. Equation (33) should therefore be a good approximation.

Because *f_LOH_* is probably insensitive to tumour size *N*, it should be possible to estimate *n_X_* from experimental *f_LOH_* figures even when the sizes of the sampled tumours are unknown or have been lost. Equation (33) implies:

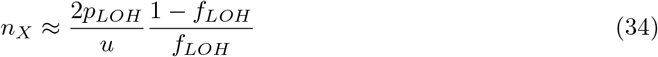

Without an LOH survey of malignant schwannoma samples, this formula can’t yet be used to provide a new estimate of *n_X_*. It is presented here for future work.

Returning to the second point about whether or not this experiment would be realistic, we should now estimate *f_LOH_*. Substituting the estimates *p_LOH_* = 7.97 × 10^−8^, *u* = 4.48 × 10^−10^ and *n_X_* ≈ 1245 into equation (33) yields

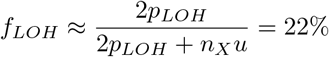

This is a large enough value that a pilot study with only 15 samples is likely to detect something: the probability that *no* samples display excess LOH, assuming the above estimates, is (1 −0.22)^15^ = 2.4%.

It is notable that (33) is independent of tumour size N. A study of biopsies from malignant VS tumours should therefore not detect an association between LOH and tumour size. It should be remembered that this assumes selective neutrality of *s_l_* and *s_m_* – i.e. *TSX* is haplosufficient.

## 4 Modelling excess risk of malignancy following irradiation

The main mechanism of mutagenesis following radiation is the induction and misrepair of doublestrand breaks (DSBs) [50, 51, 52, 53]. Our model of radiation mutagenesis can be summarised in simple terms: following a large dose of radiation *D*, a DSB occurs at a given base pair with some small probability *p_DSB_* (*D*). The DSB is then repaired, with a probability *ϵ* of faulty repair. There are *m_X_* such sites on gene *TSX* where misrepair results in *TSX* being deactivated. If a DSB occurs on a gene, *and* the repair was faulty, *and* it was on one of the sensitive sites, then one copy of *X* will be lost. These are taken to be independent events. The probability *P*(*X* lost) to lose one copy of *X* in a given cell is therefore

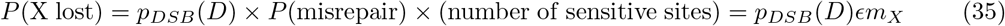

The repair error rate *ϵ*, probability of DSB induction *p_DSB_*, and number of radiosensitive sites *m_X_* now need to be specified.

There are two main mechanisms of repair following irradiation and the induction of doublestrand breaks. These are non-homologous end joining (NHEJ), and homologous repair (HR) [51, 52, 53]. NHEJ is thought to have a relatively low error rate, and HR a somewhat higher one [51]. In addition to HR, there are several other repair mechanisms with still higher error rates that take over when HR is suppressed [50]. At low dose rates, NHEJ is supposed to be the primary repair mechanism, with HR and minor mechanisms taking over at higher dose rates [50, 51, 54]. Misrepair is much more common at large dose rates of radiation, in excess of 0.5 cGy/min [55, 56, 54].

DSB misrepair is associated with the introduction of small insertions and deletions (“indels”) [51]. To model the number of sensitive sites, we will use the parameter *m_X_* from section 2, because this was the number of indel-sensitive locations. From (9), *m_X_* ≈ 0.74l_*X*_, or in terms of *n_X_*, the total number of sensitive sites on gene *X*,

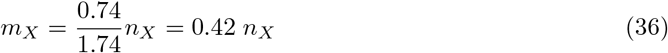

The therapies we are modelling have high dose rates, on the order of several Grays per minute: this is far outside the low dose rate regime studied by Stenerlöw et al. [56]. The repair in this regime will therefore be relatively error-prone, similar to that studied by Rothkamm et al. In this high dose rate regime, as many as 50% of the repairs may be faulty [55]. Strictly speaking, the error rate should depend on both dose *D* and dose rate 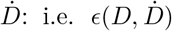. In the absence of sufficiently well-developed models, we will instead take *ϵ* = 50% = constant [55], and leave more detailed models of DSB repair to future work. The figure of *ϵ* = 50% is a pessimistic upper bound: this section should then establish an upper limit on the associated risk.

The only term in the above model that has not been fixed is *p_DSB_*(*D*), the probability for a DSB to be induced. This has been measured directly, at the same energies and doses of gamma rays that are used in therapy [57]. In the 0 – 50 Gray regime, the results are well described by a simple linear interpolation,

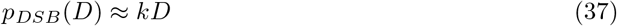

with *k* = 3.90 × 10^−7^ per Gray per base pair. Over the entire genome, this corresponds to about 300 induced DSBs per Gray of radiation. This is substantially higher than older measurements of mutagenesis using X-rays – this is consistent with the understanding that gamma radiation is more mutagenic [56, 54, 57].

To result in malignancy, the cell carrying the new mutation has to survive. Following a dose of ionizing radiation, a fraction of any population of cells will fail to replicate, and die. We will assume that all cells in the tumour have the same probability *S*(*D*) to survive the dose *D* administered. Following current models in radiobiology, we will use a “linear-quadratic” form for *S*(*D*) [58, 59]:

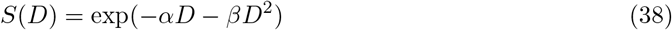

and take *α* = 0.77Gy^−1^ and *β* = 0.31Gy^−2^ [58, 59].

Combining equations (35), (37), and (38), the probability that after a dose of radiation, a given cell loses one copy of *TSX and survives* is

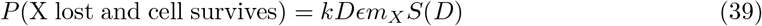

Because *kD* will be on the order of 10^−5^, and *ϵ* and *S* both on the order of 1, we can only expect this probability to be very large when *m_X_* ≈ 10^5^. The largest known gene in the human genome, dystrophin, is 2.5 Mb long – the majority are much smaller [60]. It should be safe to assume that any candidate *TSX* will be short enough that the probability (39) is much smaller than 1.

Not all cells will become cancerous when hit by a dose of radiation. Cells that still have two copies of tumour suppressor *X* can lose one copy and still retain one functional copy. Cells that have already lost one copy of gene *X* may become malignant if they lose the other. The only subpopulations that will be sensitive to irradiation are therefore *N_l_* and *N_m_* from figure 8.

The probability that a tumour gains a new malignant clone following irradiation should therefore be:

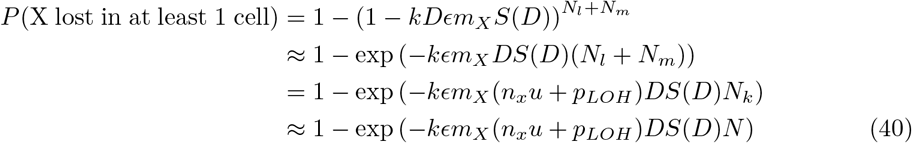

where we have used the solutions (24) and the approximation *N_k_* ≈ *N*. The total number of cells in the tumour *N* can again be related to the volume of the tumour using equation (28).

Equation (40) gives the probability that the gene *TSX* will be lost. This can only result in malignancy if it has not been lost already. The probability to have developed a malignancy after a dose *D* should be

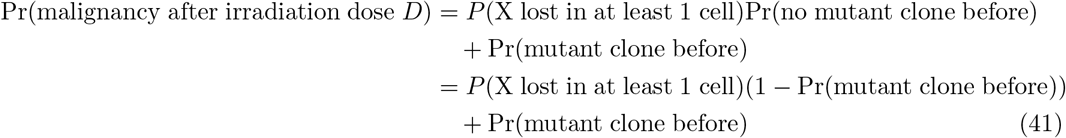

i.e. there is some probability P(mutant clone before) that an undetected malignancy already existed.

Current practice defines the *excess* risk *E.R*. of malignant transformation as the *absolute risk difference* [61]: the difference between the probability Pr(malignancy after irradiation dose *D*) and the probability of a malignancy having occurred spontaneously beforehand, Pr(mutant clone before):

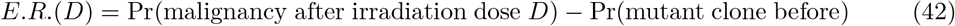

From equations (42) and (41), it should be clear that

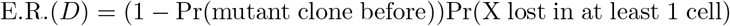

and finally, substituting equation (26) for Pr(mutant clone before) into the above yields the main result,

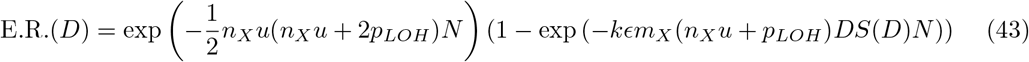

Some initial observations can be made. *E.R*.(*D*) initially increases linearly in dose *D*, with no lower threshold. At some point, the exponential fall-off of *S*(*D*) will cause E.R. to peak, then decrease rapidly. The *stochastic* effect of radiation-induced mutations is eventually overcome by the *deterministic* effect of necrotic cell death. It is remarkable that the linear dependence at low doses, below around 2 Gray, is consistent with the linear no threshold model [58].

### 4.1 Dose fractionation

Equation (43) assumes that the dose *D* is delivered in a single fraction. However, it is more common to deliver treatments in multiple fractions. This has been demonstrated to provide effective tumour control whilst also reducing side effects in healthy tissue [62, 63]. Fractionation schedules need to be chosen in order to balance what are typically referred to as the “5 R’s of radiotherapy”: repopulation, repair, redistribution, reoxygenation and radiosensitivity. Fractionation is beneficial from the perspective of tumour control, as it allows for the redistribution and reoxygenation of tumour cells during the course of treatment, whilst also allowing healthy cells a chance to repair and repopulate.

Fractionated doses should be administered so that tumour cells have no or little time to recover. Delivered in rapid enough succession, the *total* dose *D* will be what matters, and a tiny proportion of tumour cells will survive due to the rapid fall-off in *S*(*D*). How closely doses need to be spaced to prevent tumour cell recovery likely depends on the cell-cycle time. This is the rationale behind scheduling fractions to be delivered every day, or every two days.

A typical cell cycle time for Schwann precursor cells is not known to a high degree of precision. The *b* = 25.5/yr estimated in section 2.1 corresponds to a cell cycle time of around 14 days, but this was for wild-type precursor cells, not tumour cells. We can estimate the cell cycle time for tumour cells using tumour expansion speed c and Schwann cell volume *V_SC_* via a scaling argument:

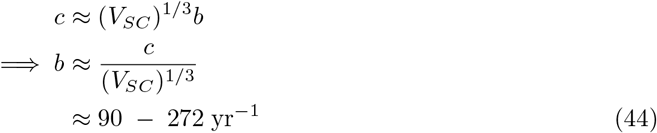

with *V_SC_* = 1.6 × 10^−6^mm^3^ from [45], as in equation (28); and *c* =1 – 3mm/yr from [64]. This rough estimate should be consistent with estimates from both Fisher-Kolmogorov and Eden model approaches to solid tumour growth [65, 66]. Note that this estimate is faster than the precursor cell division rate, which is consistent with tumour cells having a small selective advantage.

The surface growth rate *b* estimated in this way corresponds to a cell division timescale of 1 – 4 days. As long as fractionated doses are spaced more closely than this, they should have the same effect on tumour cells as a single large dose, and the excess risk will be governed by equation (43), and not equation (46), below.

However, enough is unknown about the dynamics of Schwann cells *in vivo* that we should also consider a pessimistic scenario, in which tumour cells manage to fully recover between fractionated doses. This should give us an upper bound – the worst case scenario – on the risk of properly fractionated therapy, with (43) giving the lower bound – the best case scenario.

In this worst case scenario, we treat the risk of mutations being induced following each fractional dose as *completely independent events*:

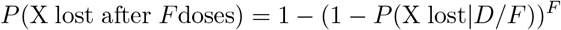

which implies

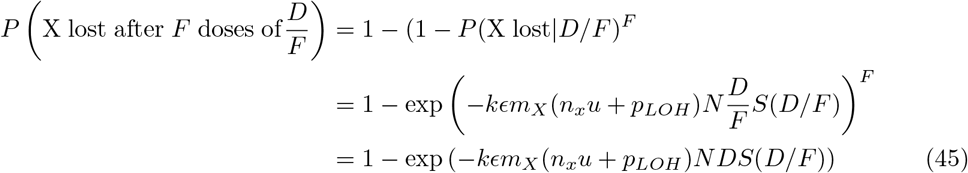

and the excess risk associated with a dose *D* delivered in *F* fractions is

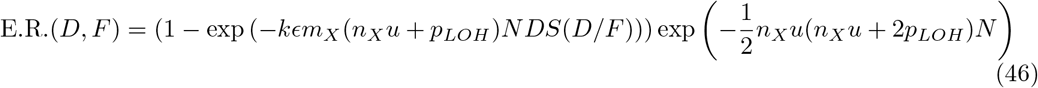

The likelihood of inducing a new mutation is essentially unchanged, but the likelihood of *survival S*(*D/F*) now depends strongly on the number of fractions *F*. This has important implications in section 5.3.

## 5 Results

### 5.1 Sporadic vestibular schwannoma

The answer to our first research question, on modelling the risk of benign vestibular schwannoma, is that observed incidence can be explained well with a mechanistic, three-hit model involving *NF2, SMARCB1*, and one additional oncogene. The simple model in section 2 has a clear interpretation for each of its parameters, which allowed us to find new estimates for the rates *u* of point mutations and *r_LOH_* of LOH, which improve on current estimates for Schwann cells (see figure 3). The technique of estimating parameters and bootstrapping uncertainties from measurements of the relative frequencies *f_LOH_* and *f_SMARCB1_*, derived from experimental data, should also be applicable to other neoplasias (see section 2.2 and figure 3).

The estimation technique for *rLOH* is very sensitive to the experimental frequency *f_LOH_* of LOH events. If an experiment can’t detect all forms of LOH present, *r_LOH_* will be underestimated. We have tried to anticipate this by drawing on studies that explicitly account for copy-neutral LOH [25]. Nonetheless, it is natural to ask why our estimate of *r_LOH_* is almost 70 times lower than a previous estimate for colorectal cancer [10]. First, the cell division rate *b* is much slower in glia (*b* = 25.5/yr [23]) than in the colonic crypt (every 5d implies *b* = 73/yr [67]), so the values of *r_LOH_* cannot be directly compared. We should instead compare *p_LOH_* = *r_LOH_*/*b*, the probability of an LOH event per division, and account for uncertainties in both estimates of *p_LOH_*.

Performing a similar bootstrapping procedure for the estimates of *p_LOH_* for colorectal crypt cells by resampling data from Huang et al. 1996 and using the same calculation as Paterson, Bozic and Clevers 2020 to determine *p_LOH_* confirms that there is still a statistically significant difference – a factor of 23 (see figure 10) [68, 10]. It may be the case that rates of LOH are simply different between glial cells and colonic crypt cells, in the same way that point mutation rates are known to vary substantially between tissues [28, 67]. P. Herrero-Jimenez et al. reported that *p_LOH_* may vary by a factor of 30 between colonic crypt and hematopoietic stem cells, so a difference of a factor of 23 between different tissues is quite possible [69].

The number of sensitive sites on the third hit, *n*_3_, was found to be 2002. This high value may indicate that there are several oncogenes that contribute to sporadic incidence, representing a high level of genetic diversity. This is also likely to be an overestimate of the real figure, and may be revised once *LZTR1* is taken into account.

The posterior distribution for *n*_3_ was found to be bimodal, with two peaks around 700 and 2000 (see figure 7). We believe this is due to the low rate of *SMARCB1*-positive samples in our experiments, which implies a high uncertainty in equation (13). A larger sample size for *SMARCB1* levels would be very helpful to constrain n_3_ further.

It is also interesting to note that the theory developed in section 2 predicts that there should be no association between the relative frequency *f_LOH_* of LOH and patient age. The same holds for *f_SMARCB1_* This is a consequence of neutrality: a change in allele frequency over time would indicate selection for one of the intermediate genotypes in figure 2 [70]. Previous work also suggests that indirect knowledge of selective advantages may be contained in the orders of appearance of mutations, even when the full dynamics over time are not available [10].

#### 5.2 Malignant transformation

Our second goal was to establish the risk of sporadic malignant transformation in an untreated schwannoma. The model in section 3 culminates in equation (29): a simple linear relationship between risk of malignancy and tumour volume *V*. The most interesting finding here is that most of the observed lifetime risk of malignancy can be explained by sporadic incidence (see figure 11) [2]. If malignant transformation is associated with the loss of an unidentified tumour suppressor, then plausible values for *n_X_* and tumour volume can explain much of the observed lifetime risk (see figure 11).

This should be qualified by the uncertainty involved. The tumour suppressor TSX is currently unidentified, so a wide range of estimates for risk can be produced by adjusting *n_X_*. Furthermore, our early estimates of *n_X_* suggest a value on the order of 1245, which while not impossible is quite high. This may suggest that there is a degree of genetic diversity that our modelling approach has not yet captured. On the other hand, there are tumour suppressors implicated in other cancers known to have higher *n_gene_* values, so there is some chance that one gene explains most of the incidence [10].

To try and locate the genes involved in malignancy, we sketch an experimental study in section 3.2. Our calculations suggest that “excess” LOH should be a frequent enough occurrence to detect by profiling at least 15 tumours. This might be achievable by looking for copy number changes – but a higher resolution approach like an SNP array would be more sensitive to copy-neutral LOH and smaller deletions [71, 72]. Although this is not a detailed experimental design, this at least suggests that such an experiment should be feasible.

As was the case with the model of sporadic incidence, these levels of LOH should not be associated with size in any significant way. This is a consequence of neutrality: an association with size would indicate selection for one of the intermediate mutants [70], and thus point to haploinsufficiency of the implicated gene. However, our neutral model is not detailed enough to comment on the magnitude or direction of this effect.

#### 5.3 Excess risk of radiotherapy

Our final goal was to develop an *a priori* model of the excess risk of malignant transformation following irradiation. For both unfractionated radiosurgery and for fractionated radiotherapy under ideal conditions, the excess risk of malignancy was found to be totally negligible for realistic doses. As can be seen in figure 12, the excess risk of malignant transformation for a total dose of > 10 Gray is remote, less than 1 in a billion. This should hold for a wide range of tumour sizes and values of *n_X_*. This is consistent with previous studies that found radiotherapy had no detectable association with new malignancies [3, 73].

**Figure 12:**
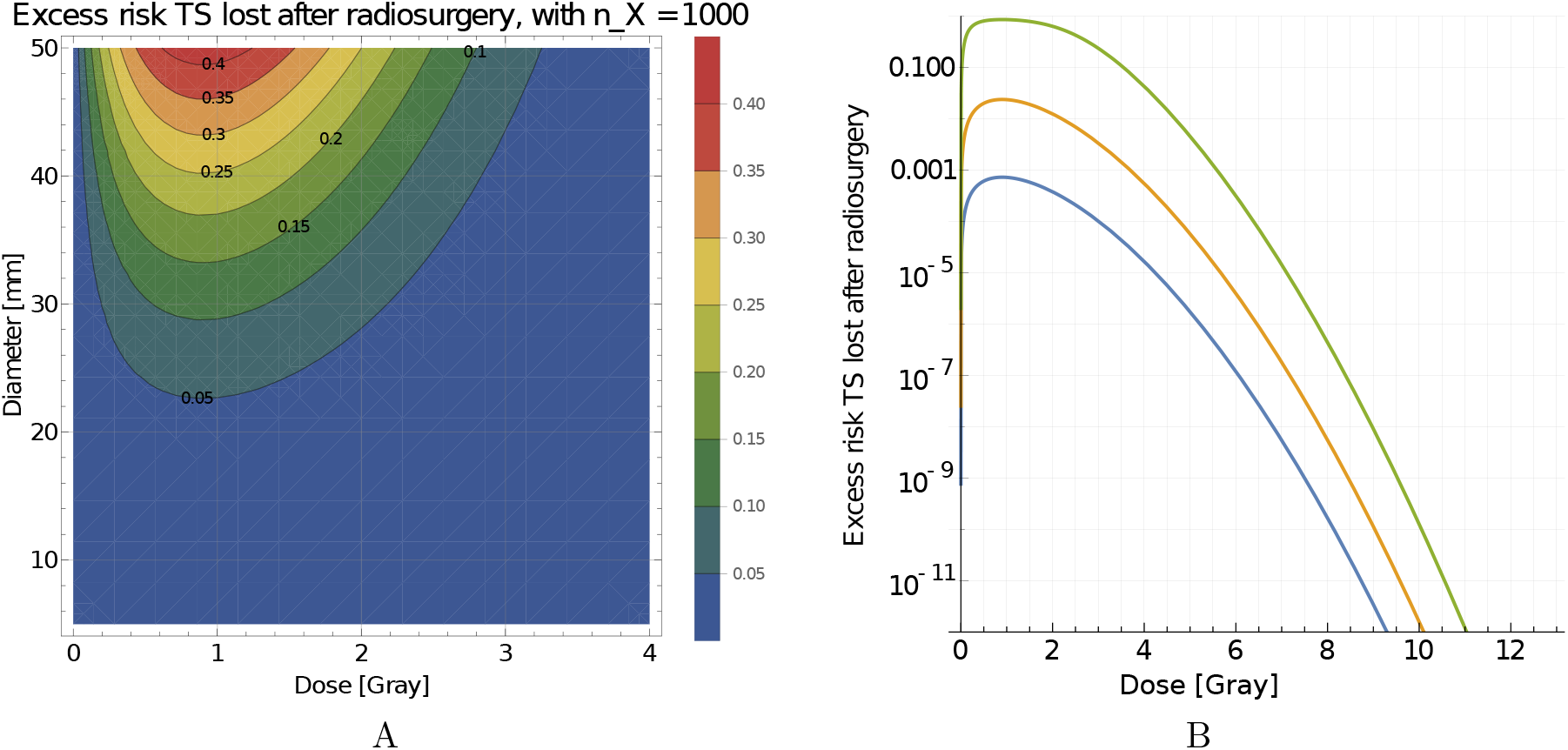
Our model for the excess risk of malignancy associated with irradiation, from equation (43). This model is based on the frequency of DSB induction by irradiation, and accounts for both tumour volume and cell survival (see section 4) [57, 58, 6]. The dose is assumed to be delivered in a single fraction, as in stereotactic radiosurgery. Fractionated radiotherapy should have a similar dose-response relationship if scheduled properly (see section 4.1). A: The excess risk of a radiation-induced DSB deactivating a tumour suppressor as a function of both tumour diameter (*y*-axis) and dose (*x*-axis), for *n_X_* = 1245 (best estimate from section 3.1) and *m_X_* = 0.42 × 1245 = 523 (from equation (36)). B: The risk of deactivating a tumour suppressor in a tumour of diameter 20 mm as a function of dose: for values of *n_X_* of 100 (lower curve, blue), 1000 (middle curve, yellow), and 10^4^ (upper curve, green), and values of *m_X_* = 0.42*n_X_* (equation (36)).

For lower doses, on the order of 1 – 4 Gray, the risk is much greater. These doses are much lower than therapeutic targets, so current best practice should have a minimal risk. However, for cases where there are many diffuse tumours in the same region, it may not be possible to target individual tumours with optimal dosing. The relative risk for these cases should be revisited as a priority, ideally combining the above model with medical imaging data.

It must be emphasised that our conclusions about radiotherapy are theoretical, and an indirect extrapolation from experiments and incidence studies. We have tried to account for the inevitable uncertainty in this by erring on the side of pessimism: for example, with regards to the DSB repair error rate *ϵ* in section 4. This is so that we can estimate a reasonable upper bound on risk, despite the unknowns.

If doses are spaced so that fewer than one cell cycle passes for the tumour cells, so that the cells “see” the fractionated doses as a single, large dose, then the risk of malignancy should be negligible. We estimate the cell division time-scale to be on the order of 1 – 4 days (section 4.1). If at least 4 Gray can be delivered each cell cycle, the excess risk of malignancy should be less than the expected lifetime risk, even for pessimistic estimates of *n_X_* (see figure 12).

Slow-growing schwannomas can probably be treated safely at a minimum dose rate of 1 Gray per day. Faster growing tumours could justify more aggressive treatment: the 3mm/yr cases from Paldor et al. 2016 might require 4 Gray per day to ensure that we are on the right side of the peak risk from figure 12 [64]. It would be interesting to learn how this difference in growth rates is correlated with specific genetic alterations.

This does assume that solid growth can be reliably distinguished from inflammation and “pseudo-progression”, which may not always be possible.

For completeness, we also studied the “worst case scenario” in which fractionated therapy is delivered in independent doses several days apart. This would maximise the chance that tumour cells “recover” from the deterministic effects of cell death, and go on to divide again. This should not be the case in practice [62].

As the number of fractions increases, the proportion of mutant cells that survive the therapy also increases. As a result, poorly administered hyperfractionated therapy shows a much higher risk for realistic doses and fractions. This risk overtakes the lifetime risk at *F* ≈ 10 fractions, and may grow as high as ≈ 10% (see figure 13). This enhancement of risk is attributable to the increased survival of mutants following therapy, and not due to enhanced mutagenicity as such (see figure 13).

**Figure 13:**
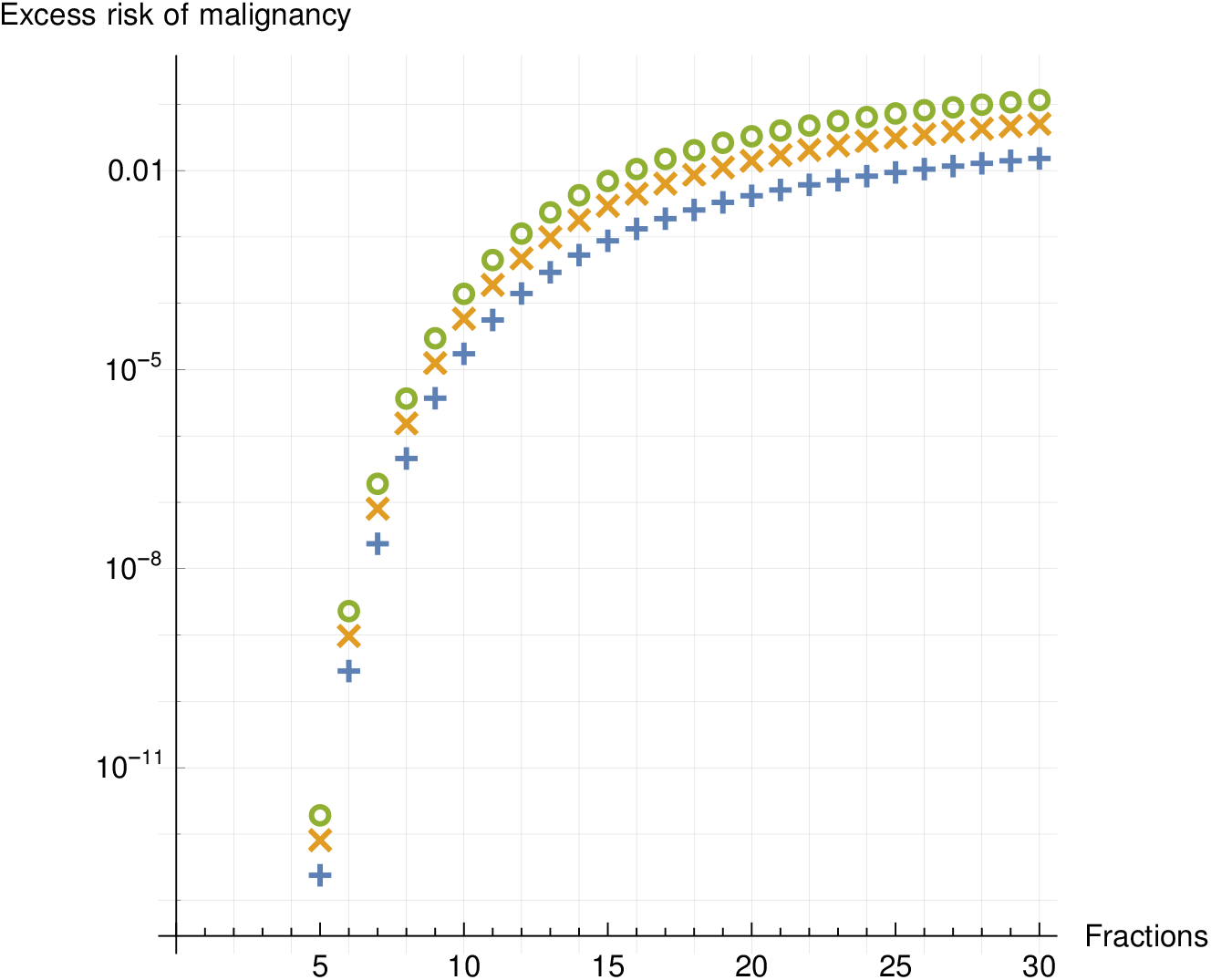
Worst-case scenario excess risk of radiation-associated malignant transformation for tumours of diameters 20mm (+ signs), 30mm (× signs), and 40mm (○ signs) each receiving a dose of 50 Gray, if the two hits are a hypothetical gene with *m_X_* = 100. The dose is delivered in separate fractions at least a week apart, maximizing tumour cell recovery. There is a very sharp dependence on the number of fractions the dose is split into (*x*-axis). This sharp dependence is attributable to the way that the cell survival curve *S*(*D*) falls off in equation (40).

These results must be qualified by the uncertainties in the identity of the hypothetical tumour suppressor (or suppressors), and also in the dynamics of cell division, DNA repair and survival. In addition, our model for the cell survival curves *S*(*D*) could be improved upon. This may effect on our conclusions regarding the “worst-case scenario”, but probably not best practice.

## 6 Discussion

In addition to improved estimates of the underlying mutation rates and dynamics of LOH in Schwann cells, our mechanistic approach to modelling schwannomas has enabled new theoretical estimates of the excess risk of malignant transformation in radiosurgery. The uncertainties in this model have raised some interesting questions, and suggested new opportunities for future modelling and experimental work.

Firstly, a better picture of genetic diversity in sporadic VS may be achievable with a more detailed model. This more detailed mechanistic model could include *LZTR1* as one of the first hits, as well as explicitly studying candidate oncogenes as third hits. This could be coupled with a comprehensive study of LOH in VS biopsies, as well as the status of *SMARCB1, NF2, LZTR1*, and candidate oncogenes. This might be achieved by coupling a copy number analysis with an SNP array or a similar method to detect copy-neutral LOH [71, 72]. A more complex model would require new experiments to determine its parameters, but the method for doing so should be essentially similar to the method in section 2.1: for each gene in the new model, count the total number of cases where pathogenic variant alleles were detected, and compare the allele frequencies *f_gene_* to those predicted by the model.

When applied to malignant transformation, our approach showed that the observed lifetime risk can be explained by a simple two-hit model in the growing tumour (see figure 11). This theory also predicts excess LOH on the locus of the tumour suppressor (or suppressors) responsible, *TSX*. Our estimates of n_X_ in section 3.1 suggest a promising experimental approach would be to look for LOH in at least 10 samples of malignant schwannomas. This might uncover new risk factors, as well as markers of malignant transformation.

The extension of this modelling approach to radiotherapy suggested that at tumour sizes and radiation doses typical of therapy, the excess risk of malignancy for sporadic schwannoma is negligible. Cells that receive high doses, and thus have the highest likelihood of induced mutations, will be the least likely to survive. Stochastic effects, which include malignancy, should peak at intermediate doses of around 1 Gray. Above about 4 Gray, deterministic effects dominate, and cell survival falls off very rapidly. This dose is well below recommended prescriptions [63, 74].

The excess risk should also be very small for fractionated therapies, as long as dose rates are higher than 4 Gray over the course of one tumour cell cycle, which we estimate at around 4 days (see section 4.1). This is broadly in line with current recommendations. Current clinical practice is therefore expected to have a negligible excess risk of radiation-induced malignancy, consistent with previous empirical studies [7, 3, 73].

The highly pessimistic worst-case scenario in which tumour cells fully recover in between doses should be completely avoidable if the most rapidly growing tumours are treated more aggressively (3mm/year expansion = 4 Gray/day) than more common cases. This pessimistic scenario, detailed in section 4.1, is unlikely. Nonetheless, it may be interesting to investigate whether there is an association between malignancy and failure to complete treatment. It should also be cautioned that we do not consider cases of familial *NF2*, and furthermore assume that tumours are well-isolated and recieve homogenous doses.

These findings need to be qualified by the uncertainties both in the identity of the relevant genes, and in the mechanisms that cause death of tumour cells. These are questions for which experimental answers are urgently needed. As well as revealing new connections between genomic studies and microscopic mechanisms, this work underscores the critical need to identify the genes responsible for both tumour initiation and malignant transformation.

## 7 Acknowledgements

The authors are grateful to everyone who contributed comments and constructive criticism to the unfinished manuscript. Special thanks are due to Philippa Paterson, who proofread the manuscript extensively and verified equations (13) and (14); to Joshua D Hellier for constructive discussions regarding our statistical methods; and to Mohammad Obeidat for confirming that equation (37) and corresponding value for *k* were consistent with his experiments.

## A Series solution of sporadic incidence model

The system of differential equations (1) is linear, and solutions can be found by a variety of methods. Exact symbolic solutions are not very useful in our case. The evolutionary dynamics are neutral, with no exponential clonal expansion during initiation. Furthermore, we are interested only in relatively early times *t* compared to 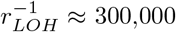 years. Our initial conditions are also not on a singular point [75, 76]. These facts make a power series solution appropriate. This also makes the similarity of the three-hit model to the classic Armitage-Doll curve very clear [4, 21].

Write

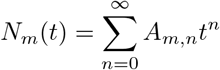

for subpopulation *m* (see figure 2 and system (1)). The *A_m,n_* coefficients can then be found recursively [75].

The resulting solutions, truncated at the third term, read:

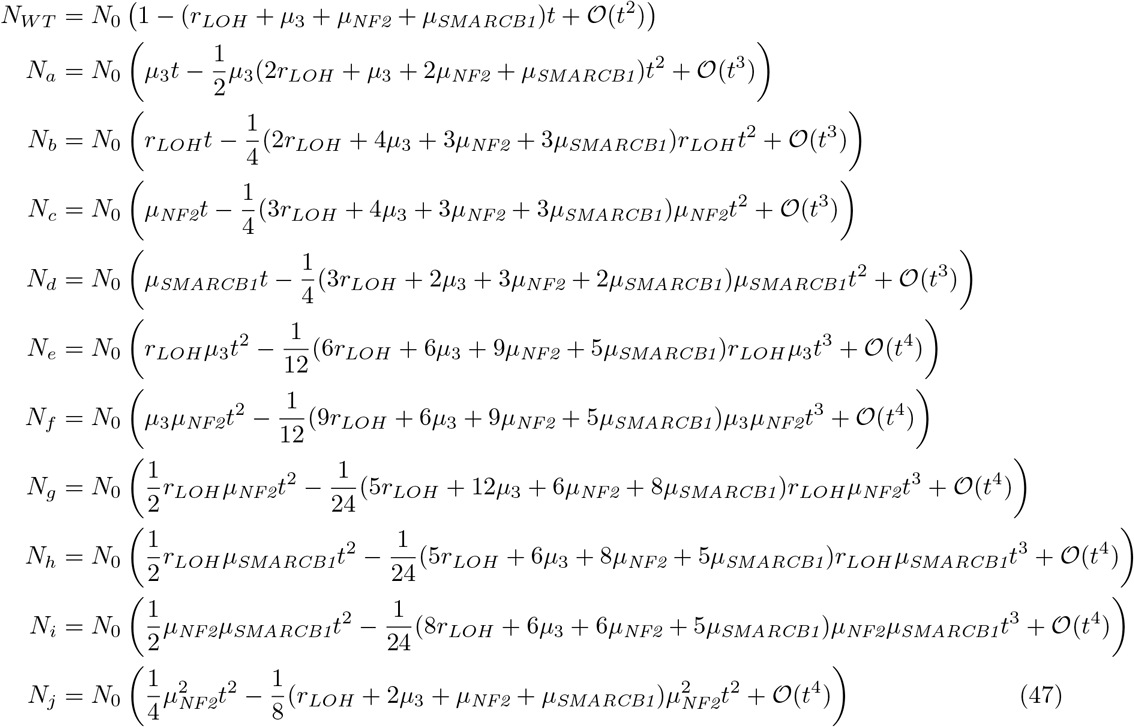

and so, *P*_1_, *P*_2_ and *P*_3_ read

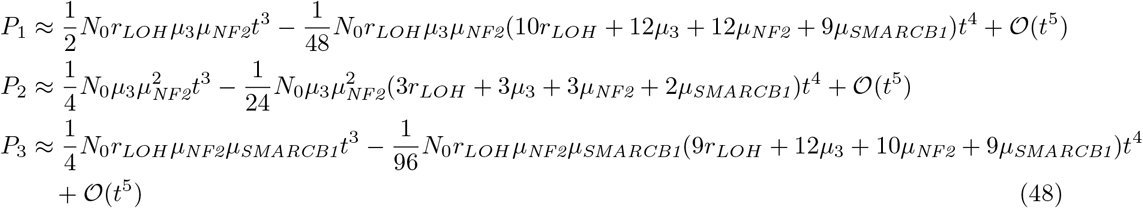

It is also clear that *f_LOH_* and *f_SMARCB1_* are only approximately independent of patient age:

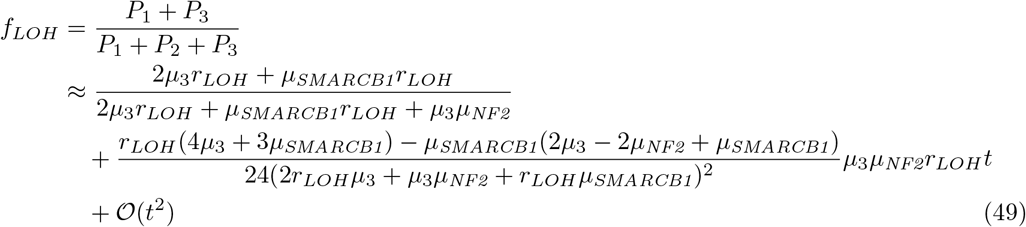

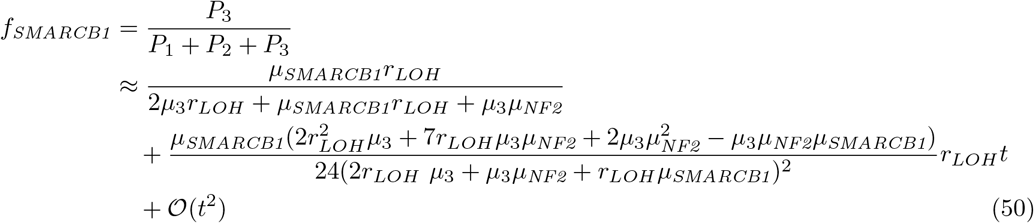

It follows that this approximation is accurate up to a factor of approximately 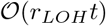, so any age-dependence should only show up on a timescale of 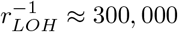 years. This is far in excess of a typical human lifetime, so in any real empirical study, any association can be expected to be unobservable.

This assumes that these alterations are selectively neutral. If this is not true, then the relative frequency of these alterations may change over time, indicating selection [70]. It may also be the case that the model ceases to be accurate over shorter timescales than 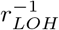: without explicitly developing and solving a non-neutral model, we cannot comment further.

## B Statistical methods: bootstrapping and regularisation

Bootstrapping consists of drawing new samples at random from a dataset, with replacement [41, 39]. Our measures of relative frequency come from datasets with *n* samples, *k* of which are positive results. When one sample is drawn uniformly at random, there is a probability *p* that this draw will be positive.

Bootstrapping n samples with this probability *p* will result in a “sample” value 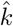. Because this sampling process consists of *n* independent draws with replacement, the resulting sample value 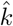 will be binomially distributed:

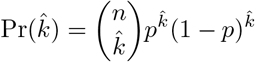

These draws can therefore be *simulated* by generating binomially distributed random numbers, given an estimate of the parameter *p*.

Binomial random variables were generated with Python’s NumPy library (https://numpy.org/about/). For each sample value 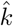, a sample frequency estimator 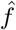 can be calculated:

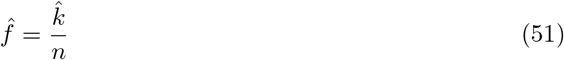

and substituted for *f_LOH_* or *f_SMARCB1_* in formulae (13) and (14). Repeating this process for many values of 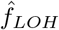 and 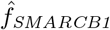 allowed us (in theory) to generate distributions for *n*_3_ and *r_LOH_* that reflect the uncertainty due to the small sample size of the underlying experiment.

However, a problem arises if 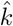 is 0 or *n*: when 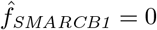 or 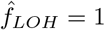, equations (13) and (14) are singular. This can make distribution means, variances, and even some quantiles ill-defined. To address this, we used a Bayesian estimator for 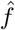 given 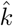 and *n* [36, 34],

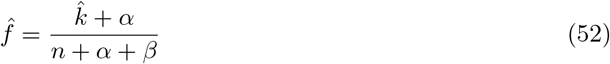

for some *α* = *β* > 0. The exact choice of constant *α* = *β* was not found to have a strong effect on our estimates or computed confidence intervals: we chose the Jeffreys prior, 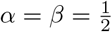, so that

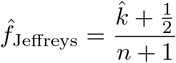

This gives a rigorous justification for additive smoothing: it ensures that our prior distribution is uninformative, and regularises the sample estimator 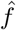, making it well-behaved when 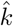 is 0 or *n* [34]. Intuitively, this can be thought of as adding one additional datum with a value of 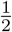 to the resampled dataset.

It should be noted that we already apply additive smoothing with 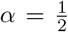 to our estimate of the parameter *p* in 2.1.3 before the bootstrapping procedure. This avoids the case in which *p* = 0, in which case the results would be meaningless: it is impossible to meaningfully resample data with no events. This is therefore not a true bootstrapping procedure, in which we resample the underlying data. It is instead a simulation of a bootstrapping procedure applied to a regularised dataset. Namely, this regularised dataset is equivalent to adding one extra datum to the original, with a value of 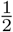.

It is easily seen that this procedure becomes equivalent to bootstrapping in the limit that *α* = 0.

We also considered and rejected the use of an informative prior. Because the regularisation procedure is applied both before and after resampling, an uninformative prior should be used in both cases, so as not to introduce inappropriate bias. From a Bayesian perspective, while there are good reasons to believe that *f_LOH_* and *f_SMARCB1_* are neither exactly zero nor exactly one, there is no objective reason to pick a particular value otherwise.

The choice of prior is easily adjusted to see what effect it has on our conclusions: our computed confidence intervals are relatively robust. The Python code that implements the bootstrapping procedure has been included in the supplemental material.

