## supplemental_code for "A mechanistic mathematical model of initiation and malignant transformation in sporadic vestibular schwannoma": bootstrap.py

```
#!/usr/bin/python3

from numpy.random import binomial
import numpy as np
from matplotlib import pyplot as plt

import seaborn as sns
import pandas as pd
import sys

# Gene sensitivity coefficients:
nNF2 = 135
nSMARCB1 = 85

def beta_est(p, n, k):
    # test: bayesian estimator with INFORMATIVE beta prior
    # we use a beta destribution as a prior, and choose alpha and beta so that
    # the expected value of k/n = p, and the probability that k==n or k==0 is
    # zero.
    if (p > 0.5):
        a = 2 * p / (1 - p)
        b = 2
        return (k + a) / (n + a + b)
    if (p < 0.5):
        a = 2
        b = 2 * (1 - p) / p
        return (k + a) / (n + a + b) 
    else:
        return (k + 2) / (n + 4)

def unbiased_est(n, k):
    # frequentist estimator, but regularised so things don't blow up.
    reg = 0.1 / n
    return (k + reg) / (n + reg) # no x2 on the bottom

def jeffreys_est(n, k):
    # uses a beta distribution prior with alpha = beta = 0.5 (Jeffrey's prior)
    return (k + 0.5) / (n + 1.0)

def bayes_est(n, k):
    # uses a beta distribution prior with alpha = beta = 1
    return (k + 1.0) / (n + 2.0)

def squashy_est(n, k):
    # uses a beta distribution prior with alpha = beta = 2 (uninformative
    # prior, avoids extremes)
    return (k + 2.0) / (n + 4.0)

def haldane_est(n, k):
    # approximates a haldane prior
    epsilon = 0.1
    return (k + epsilon) / (n + 2 * epsilon)

def sample_f(n, kobs):
    p = float(kobs) / n
    k = binomial(n, p)
    return jeffreys_est(n, k)

def n3(floh, fsmarcb1):
    nSMARCB1 = 85

    return 0.5 * nSMARCB1 * (floh - fsmarcb1) / fsmarcb1

def ploh(floh, fsmarcb1):
    # this is rloh / u b
    if (floh > fsmarcb1):
        return 0.5 * nNF2 * (floh - fsmarcb1) / (1 - floh)
    else:
        return 0

def plohCRC(f):
    nAPC = 604
    u = 1.70*10**(-10)
    k = np.sqrt(1.0 + f / (1.0 - f)) - 1
    return 10 * k * nAPC * u

def u(A, n3, ploh):
    b = 25.5
    N0 = 450E3

    Aobs = 0.25 * N0 * nNF2
    Aobs = Aobs * (n3 * (2 * ploh + nNF2) + nSMARCB1 * ploh)
    X = A / Aobs
    X = X ** (1. / 3)
    u = X / b

    return u

def nX(N, C, ploh, u):
    if (ploh > 0):
        B = (C * 2.0) / (N * ploh * ploh)
        nx = ploh * (np.sqrt(1.0 + B) - 1.0) / u
    else:
        nx = np.sqrt(B)/u
    return nx

def mean(varbl):
    mean = 0.
    for x in varbl:
        mean += x
    mean /= len(varbl)
    return mean

def var(varbl):
    X = 0.
    for x in varbl:
        X += x*x
    X /= len(varbl)
    mu = mean(varbl)
    return X - mu * mu

def median(varbl):
    varbl.sort()
    alpha = 0.05
    size = len(varbl)
    median = varbl[int(0.5 * size)]
    low = varbl[int( (alpha / 2.0) * size)]
    hi  = varbl[int( (1.0 - alpha / 2.0) * size)]
    return [varbl[0], low, median, hi, varbl[-1]]

def indicate(k):
    # cute indicator:
    indi_period = 40000
    indi_switch = indi_period / 4
    if ((k % indi_period) == 0):
        print("-", end='\r')
    if ((k % indi_period) == indi_switch):
        print("\\", end='\r')
    if ((k % indi_period) == 2 * indi_switch):
        print("|", end='\r')
    if ((k % indi_period) == 3 * indi_switch):
        print("/", end='\r')

def makefigures(show,Nbins,results):
    plt.rcParams['figure.figsize'] = (4,4)

    sns.set()
    sns.set_theme(style="whitegrid")
    ix = 1
    for result in results:
        dist = pd.DataFrame(result[0], columns=['Value'])
        label = result[1]
        f, (ax_box, ax_hist) = plt.subplots(2, sharex=True, sharey=False,
                gridspec_kw={"height_ratios": (.15, .85), 
                             "hspace" : 0})

        if (label == "$r_{LOH}$"):
            plt.xlim([0,1e-5])
        if (label == "$u$"):
            plt.xlim([0,1.2e-9])
        if (label == "$p_{LOH}(VS)$"):
            plt.xlim([0,3e-7])
        if (label == "$p_{LOH}(CRC)$"):
            plt.xlim([1.0e-6,3.0e-6])

        # box plots:

        ax_box.minorticks_on()
        ax_box.grid(b=True, which='minor', color='#999999', linestyle='-', alpha=0.2)
        sns.boxplot(x = dist['Value'],
                    showfliers = False,
                    whis = [2.5, 97.5],
                    width = 0.8,
                    palette = "pastel",
                    ax = ax_box)
        sns.despine(ax=ax_box, bottom=True, top=False, right=False)
        ax_box.tick_params(labeltop=False,labelleft=False,labelbottom=False)

        # histograms:

        sns.histplot(data = dist, x = 'Value', 
                bins = Nbins, 
                stat = "probability",
                kde = False,
                kde_kws={'bw_adjust' : 4.0, 'cut' : 0},
                palette = "pastel",
                ax = ax_hist)
        sns.despine(ax=ax_hist, top=True, right=False)

        plt.subplots_adjust(hspace = 0.)

        plt.xlabel(label)
        plt.ylabel("Probability")
        plt.tight_layout()
        ax_hist.minorticks_on()
        ax_hist.grid(b=True, which='minor', color='#999999', linestyle='-', alpha=0.2)

        filename="bootdist"+str(ix)+".pdf"
        plt.savefig(filename)
        if (show):
            plt.show()
        ix += 1

def main():
    # the main routine
    show = ("--show" in sys.argv)
    latex = ("--latex" in sys.argv)

    # the experimental results:
    Nloh = 23 + 1 # carlson 2018 next gen
    kloh = 17 + 0.5
   # Nsmarcb1 = 28 # k hadfield
   # ksmarcb1 = 2
    Nsmarcb1 = 32 + 1 # us!
    ksmarcb1 = 0 + 0.5

    # Huang 1996
    kCRC = 360 # 412 - 52
    NCRC = 412 # sample size

    # fitting constant:
    #A = 2.409E-11 # ages trimmed at 75
    A = 2.260E-11 # ages trimmed at 80
    b = 25.5    # empirical constant

    Nsamples = 2500000
    Nbins = 33

    # Guesses:
    Nmvs = 1e10
    Pmvs = 0.002

    # This guarantees we report the same input parameter values in the paper
    # that we use in the calculation:
    if (latex):
        with open('estimates.tex', 'w') as texfile:
            texfile.write("% Gene sensitivities:\n")
            texfile.write("\\newcommand{\\nMerlin}{" + format(int(nNF2),"d") + "}\n")
            texfile.write("\\newcommand{\\nSMARCB}{" + format(int(nSMARCB1),"d") + "}\n")

    n3samples = []
    rlohsamples = []
    usamples = []
    nxsamples = []
    plohVSsamples = []
    plohCRCsamples = []

    if (not latex):
        print("Bootstrapping "+str(Nsamples)+" samples...")

    for k in range(Nsamples):
        flohsample = sample_f(Nloh,kloh)
        fsmarcb1sample = sample_f(Nsmarcb1,ksmarcb1)

        n3sample = n3(flohsample, fsmarcb1sample)
        plohsample = ploh(flohsample, fsmarcb1sample)
        usample = u(A, n3sample, plohsample)
        rlohsample = plohsample * b * usample

        nxsample = nX(Nmvs, Pmvs, plohsample * usample, usample)

        fCRCsample = sample_f(NCRC, kCRC)

        plohVS = plohsample * usample
        plohCRCsample = plohCRC(fCRCsample)

        n3samples.append(n3sample)
        rlohsamples.append(rlohsample)
        usamples.append(usample)
        nxsamples.append(nxsample)
        plohVSsamples.append(plohVS)
        plohCRCsamples.append(plohCRCsample)

        if (show):
            indicate(k) # cute indicator

    # store results of bootstrapping procedure for later:
    results=[[n3samples, "$n_3$"],
            [rlohsamples, "$r_{LOH}$"],
            [usamples, "$u$"],
            [nxsamples, "$n_X$"],
            [plohVSsamples, "$p_{LOH}(VS)$"],
            [plohCRCsamples, "$p_{LOH}(CRC)$"]]
    # TODO separate ploh sets for separate figure. these should be a single plot
    # with a logarithmic scale

    if (not latex):
        print("\ndone")
        print("Generating summary statistics...")

    # compute statistics:
    means = []
    sds = []
    quantiles = []
    for case in results:
        means.append([mean(case[0]), case[1]])
        sds.append([var(case[0]), case[1]])
        quantiles.append([median(case[0]), case[1]])

    if (not latex):
        print("done")
        print("-----------------")
        print("Bootstrapped quantiles with " + str(Nsamples) + " samples:")

        for case in quantiles:
            print(case[1] +": "+ str(case[0]))

        print("-----------------")
        print("Exact (unbiased) estimators:")

    n3exact = n3(kloh * 1.0 / Nloh, ksmarcb1 * 1.0 / Nsmarcb1)
    plohexact = ploh(kloh * 1.0 / Nloh, ksmarcb1 * 1.0 / Nsmarcb1)
    uexact = u(A, n3exact, plohexact)
    rlohexact = plohexact * b * uexact
    nxexact = nX(Nmvs, Pmvs, plohexact * uexact, uexact)
    plohVSexact = plohexact * uexact
    plohCRCexact = plohCRC(kCRC * 1.0 / NCRC)

    if (not latex):
        print("n3: " + str(n3exact))
        print("rloh: " + str(rlohexact))
        print("u: " + str(uexact))
        print("nx: " + str(nxexact))
        print("plohVS: " + str(plohVSexact))
        print("plohCRC: " + str(plohCRCexact))

        print("-----------------")

        print("Plotting bootstrapped distributions...")

    if (latex):
        with open('estimates.tex', 'a') as texfile:
            texfile.write("% Parameter estimates:\n")
            texfile.write("\\newcommand{\\nOnco}{" + format(int(n3exact),"d") + "}\n")
            texfile.write("\\newcommand{\\umut}{" + format(uexact,".2E") + "}\n")
            texfile.write("\\newcommand{\\rloh}{" + format(rlohexact,".2E")+ "}\n")
            texfile.write("\\newcommand{\\ploh}{" + format(plohVSexact,".2E")+ "}\n")
            texfile.write("\\newcommand{\\nTSGX}{" + format(int(nxexact),"d")+ "}\n")
            texfile.write("% Confidence/credible intervals:\n")
            texfile.write("\\newcommand{\\nOncoCILo}{" + format(int(quantiles[0][0][1]),"d")+ "}\n")
            texfile.write("\\newcommand{\\nOncoCIHi}{" + format(int(quantiles[0][0][3]),"d")+ "}\n")
            texfile.write("\\newcommand{\\rlohCILo}{" + format(quantiles[1][0][1],".2E")+ "}\n")
            texfile.write("\\newcommand{\\rlohCIHi}{" + format(quantiles[1][0][3],".2E")+ "}\n")
            texfile.write("\\newcommand{\\umutCILo}{" + format(quantiles[2][0][1],".2E")+ "}\n")
            texfile.write("\\newcommand{\\umutCIHi}{" + format(quantiles[2][0][3],".2E")+ "}\n")
            texfile.write("\\newcommand{\\nTSGXCILo}{" + format(int(quantiles[3][0][1]),"d")+ "}\n")
            texfile.write("\\newcommand{\\nTSGXCIHi}{" + format(int(quantiles[3][0][3]),"d")+ "}\n")
            texfile.write("\\newcommand{\\plohCILo}{" + format(quantiles[4][0][1],".2E")+ "}\n")
            texfile.write("\\newcommand{\\plohCIHi}{" + format(quantiles[4][0][3],".2E")+ "}\n")

    makefigures(show,Nbins,results)

if __name__ == "__main__":
    main()
```
