## supplemental_code for "A mechanistic mathematical model of initiation and malignant transformation in sporadic vestibular schwannoma": counter.py

```
#!/usr/bin/python3
# Python script to count numbers of sensitive sites: sites which result
# in a stop codon when a substitution or indel of length k occurs there.
#
# Depends on regex and Biopython
#
# Usage: ./counter.py EMBL_FILE

import sys
from Bio.Seq import Seq
import regex as re
from Bio import SeqIO

stop_codons = [["T", "A", "G"],
        ["T", "A", "A"],
        ["T", "G", "A"]]

def offsets(k):
    # the three bases around a deletion of length k that could result in this
    # codon becoming a stop codon
    list_offsets= [[k, k+1, k+2],
        [0, k+1, k+2],
        [0, 1, k+2]]
    return list_offsets

def read_embl(filename):
    record = SeqIO.read(filename, "embl")
    return record.seq

def read_fasta(filename):
    record = SeqIO.read(filename, "fasta")
    return str(record.seq)

def build_sequence():
    # read a file that contains a sequence
    print(sys.argv[1])
    with open(sys.argv[1], "r") as seq_file:
        sequence = seq_file.read()
        sequence_joined = sequence.replace("\n", "")
        sequence_joined = sequence_joined.replace(" ", "")
    while (len(sequence_joined) % 3 == 0):
        sequence_joined += "N"
    return sequence_joined

def find_orfs_with_trans(seq, trans_table, min_protein_length):
    # From the documentation:
    answer = []
    seq_len = len(seq)
    for strand, nuc in [(+1, seq), (-1, seq.reverse_complement())]:
        for frame in range(3):
            trans = str(nuc[frame:].translate(trans_table))
            trans_len = len(trans)
            aa_start = 0
            aa_end = 0
            while aa_start < trans_len:
                aa_end = trans.find("*", aa_start)
                if aa_end == -1:
                    aa_end = trans_len
                if aa_end - aa_start >= min_protein_length:
                    if strand == 1:
                        start = frame + aa_start * 3
                        end = min(seq_len, frame + aa_end * 3 + 3)
                    else:
                        start = seq_len - frame - aa_end * 3 - 3
                        end = seq_len - frame - aa_start * 3
                    answer.append((start, end, strand, seq[start:end]))
                aa_start = aa_end + 1
    answer.sort()
    return answer

def longest_open_reading_frame(sequence):
    table = 11
    min_pro_len = 100
    found = find_orfs_with_trans(sequence, table, min_pro_len)
    return found[0][-1]

def generate_mk_sequence(sequence):
    # generate a sequence m(k): the number of sites sensitive to deletions of
    # length k
    k = 1 # length of deletion
    mks = []

    for k in range(1, (len(sequence) - 3)):
        # for each deletion length k...
        mk = 0
        i = 0 # first codon
        while (i < (len(sequence) - k - 2)):
            # delete a base and see if the surrounding bases form a nonsense
            # codon
            for around in offsets(k):
                post_delet_codon = []
                post_delet_codon = [sequence[i + around[0]], 
                        sequence[i + around[1]],
                        sequence[i + around[2]]]
                stop_codon = (post_delet_codon in stop_codons)
                if stop_codon:
                    mk += 1

            i += 3

        mks.append(mk)
    return mks

def calculate_l_gene(sequence):
    # calculate the number of substitution-sensitive sites on this gene
    i = 0 # first codon
    l = 0 # counter
    while (i < (len(sequence) - 3)):
        # for each codon, count how many ways it can form a nonsense codon
        # a more elegant method is probably possible
        if (sequence[i] == "T"):
            if (sequence[i+1] == "A"):
                l += 2
            if (sequence[i+2] == "A"):
                l += 2
            if (sequence[i+1] == "G"):
                l += 1
            if (sequence[i+2] == "G"):
                l += 1
        else:
            if (sequence[i+1] == "G"):
                if (sequence[i+2] == "A"):
                    l += 1
            if (sequence[i+1] == "A"):
                if (sequence[i+2] == "A"):
                    l += 1
                if (sequence[i+2] == "G"):
                    l += 1
        i += 3
    l = l * 1.0 / 3.0
    return l

def horner_poly(p, q):
    # evaluate a polynomial using horner's rule
    # p is a list of coefficients from 0 to n
    p.reverse()
    b = 0
    for pi in p:
        b = pi + b * q
    p.reverse()
    return b

def uniform(p):
    a = 0
    for pi in p:
        a += pi
    return a/len(p)

def zipf(p, s):
    # evaluate the mean of a zipf distribution
    a = 0
    b = 0
    for n in range(0, len(p)):
        a += p[n] * 1.0 / (n + 1.0)**s
        b += 1.0 / (n + 1.0)**s
    return a/b

def main():
    if (len(sys.argv) == 1):
        print("invoke me with ./counter.py embl_file.1")
        exit()

    sequence = longest_open_reading_frame(read_embl(sys.argv[1]))
    #sequence = longest_open_reading_frame(build_sequence())
    print("Longest open reading frame:")
    print(sequence)
    mks = generate_mk_sequence(sequence)

    print("\nSensitivity coefficients:\n")
    print("n_gene = l_gene + m_gene\n")
    print("l_gene calculation:")
    l_gene = calculate_l_gene(sequence)
    print("l_gene = " + str(l_gene))

    # 22 1s, 46 total # frampton et al.
    # =47% should be 1s
    # Assume a geometric distribution and choose q = 1-0.47
    print("m_gene calculation:\n")

    print("geometric distribution:")
    q = 1 - 0.47
    for q in [1 - 0.47, 0.4, 0.7, 0.99]:
        print("q = " + str(q))
        m_gene = (1 - q) * horner_poly(mks, q) 
        # 1-q might be an underestimate of the normalisation constant?
        # check:
        #norm_const = horner_poly([1 for p in mks], q)
        #print(str(1.0 / norm_const) + " " + str(1- q)) # no its fine :)

        print("m_gene = " + str(m_gene))
        print("n_gene = "+str(l_gene + m_gene))
        print("m/l = " +str(m_gene/l_gene) + "\n")

    # compare ivana's approximation:
    print("\nIvana's rule:")
    print("m_gene ~= 0.74 x l_gene = " + str(0.74 * l_gene))
    print("n_gene = " + str(1.74 * l_gene))

    # try the same for a uniform distribution
    #print("uniform:")
    #print(uniform(mks))
    # and some zipf distributions:
   # print("\nzipf:")
   # print(zipf(mks, 1.1))
   # print(zipf(mks, 1.3))
   # print(zipf(mks, 1.5))
   # print(zipf(mks, 1.8))
   # print(zipf(mks, 2))
   # print(zipf(mks, 3))
   # print(zipf(mks, 7))

    # it would be interesting to try a zipf distribution that has the k=1 entry
    # appear with the same frequency:
    # 1/zeta(s) = 1-q
    pass

if __name__ == "__main__":
    main()
```
